# Statistical Principles Define an Open-Source Differential Analysis Workflow for Mass Spectrometry Imaging Experiments with Complex Designs

**DOI:** 10.64898/2026.04.08.717212

**Authors:** Ethan B. T. Rogers, Sai Srikanth Lakkimsetty, Kylie Ariel Bemis, Charles A. Schurman, Peggi M. Angel, Birgit Schilling, Olga Vitek

## Abstract

Mass spectrometry imaging (MSI) characterizes the spatial heterogeneity of molecular abundances in biological samples. Experiments with complex designs, involving multiple conditions and multiple samples, provide particularly useful insight into differential abundance of analytes. However, analyses of these experiments require attention to details such as signal processing, selection of regions of interest, and statistical methodology. This manuscript contributes a statistical analysis workflow for detecting differentially abundant analytes in MSI experiments with complex designs. Using a case study of histologic samples of human tibial plateaus from knees of osteoarthritis patients and cadaveric controls, as well as simulated datasets, we illustrate the impact of the analysis decisions. We illustrate the importance of signal processing and feature aggregation for preserving biological relevance and alleviating the stringency of multiple testing. We further demonstrate the importance of selecting regions of interest in ways that are compatible with differential analysis. Finally, we contrast several common statistical models for differential analysis, showcase the appropriate use of replication, and demonstrate model-based calculation of sample size for followup investigations. The discussion is accompanied by detailed recommendations and an open-source R-based implementation that can be followed by other investigations.

## Introduction

Mass spectrometry imaging (MSI) [1] characterizes the spatial distribution of biomolecules such as peptides, lipids, and metabolites in biological samples. Its applications range from exploring cellular heterogeneity [2] to discovering biomarkers of disease [3] to mapping responses to pharmaceutical interventions [4]. Each MSI run rasters pixel-sized areas of a sample surface with a laser or ion beam that desorbs and ionizes molecules. The ionized molecules from each pixel location are then analyzed by a mass spectrometer, which separates the molecules on the basis of their mass-to-charge (m/z) ratios. The intensities of the peaks for each m/z are informative of molecular abundances. By combining the mass spectra with the spatial coordinates, MSI reconstructs molecular distributions in each sample.

Increasingly, MSI experiments have complex designs that involve multiple conditions, multiple samples, and samples of heterogeneous composition. These designs provide useful insight into the differential abundance of analytes between sample regions and conditions. However, data from such experiments are notoriously challenging to analyze. The measurements are affected by systematic biological variation between conditions and sample regions, as well as by random biological variation between and within the samples. The signals in the spectra are further distorted by technological inconsistencies in sample preparation, ionization, and mass analysis, and produce missing values and outliers [5–8]. The resulting datasets are massive in size, in the order of 10 to 100GB per sample with thousands of pixels and spectral features, stored in complex, sometimes propriety, file formats [9]. As a result, MSI experiments with complex designs require advanced tools for signal processing, selection of regions of interest (ROI), and downstream analytical workflows.

The goals of analytical workflows can be loosely grouped into three categories: unsupervised class discovery (i.e., unsupervised ROI segmentation), supervised class prediction (i.e., classification of the type or the status of tissues or their segments, e.g., to discover biomarkers of disease) and supervised class comparison (i.e., differential analysis of analytes between regions of tissues or conditions). Given the complexity of MSI data and the visual nature of imaging, most analytical workflows focus on class discovery or class prediction. In the case of differential analysis, there remains a need for representative workflows that are both compatible with the fundamental principles of statistical experimental design and analysis and available in open-source implementations.

The statistical principles are key for characterizing the uncertainty associated with our conclusions and promoting reproducible research. The statistical principles relevant to differential analysis are as follows [10– 13]. *First*, define the variation of interest and ways to control the unwanted variation. The experimental design must define the type and number of replicates and the protocols of randomization, blocking, and quality control to account for technological artifacts. *Second*, upon data acquisition, processing, and quality control, define a statistical model that appropriately reflects the hierarchy of different sources of variation in the data and explicitly spells out the assumptions. *Third*, estimate the parameters of the model and check the assumptions are appropriate. *Fourth*, perform statistical inference by translating scientific questions into null hypotheses, deriving model-based data summaries of signal and noise, and comparing these summaries to cutoffs. *Finally*, use quantitative estimates from current data to estimate sample sizes needed to detect changes of interest in future followup experiments.

This manuscript contributes a workflow for differential analysis of MSI experiments with complex designs. The workflow operationalizes the statistical principle above in a series of concrete steps and implements these steps in open-source R/Bioconductor software Cardinal augmented by other compatible open-source R-based functionalities. Using a case study of histological samples of human tibial plateaus from knees from osteoarthritis patients and cadaveric controls as a motivating example, as well as simulated datasets, we illustrate the analysis choices considered at each step and their impact on biological conclusions. The discussion is accompanied by detailed recommendations and R-based markdown vignettes available a https://github.com/EBRogers/MSI-Arthritis-Vignettes and (easily viewed at https://ebrogers.github.io/MSI-Arthritis-Vignettes) that can be followed and adapted by other investigations.

## Materials and Methods

### Experimental OA dataset

Osteoarthritis (OA) causes progressive degradation of articular cartilage, bone remodeling, and joint inflammation [14]. The knee joint (Figure 1A) is most often affected by OA due in part to its high mechanical loading [15]. Paired lateral and medial tibial plateau samples consisting of both cartilage and bone were obtained from surgical discards of four male subjects with stage IV OA of the medial femorotibial joint undergoing total knee arthroplasty, as well as from four age-, sex-, and BMI-matched non-OA cadaveric controls (Figure 1B).

**Figure 1:**
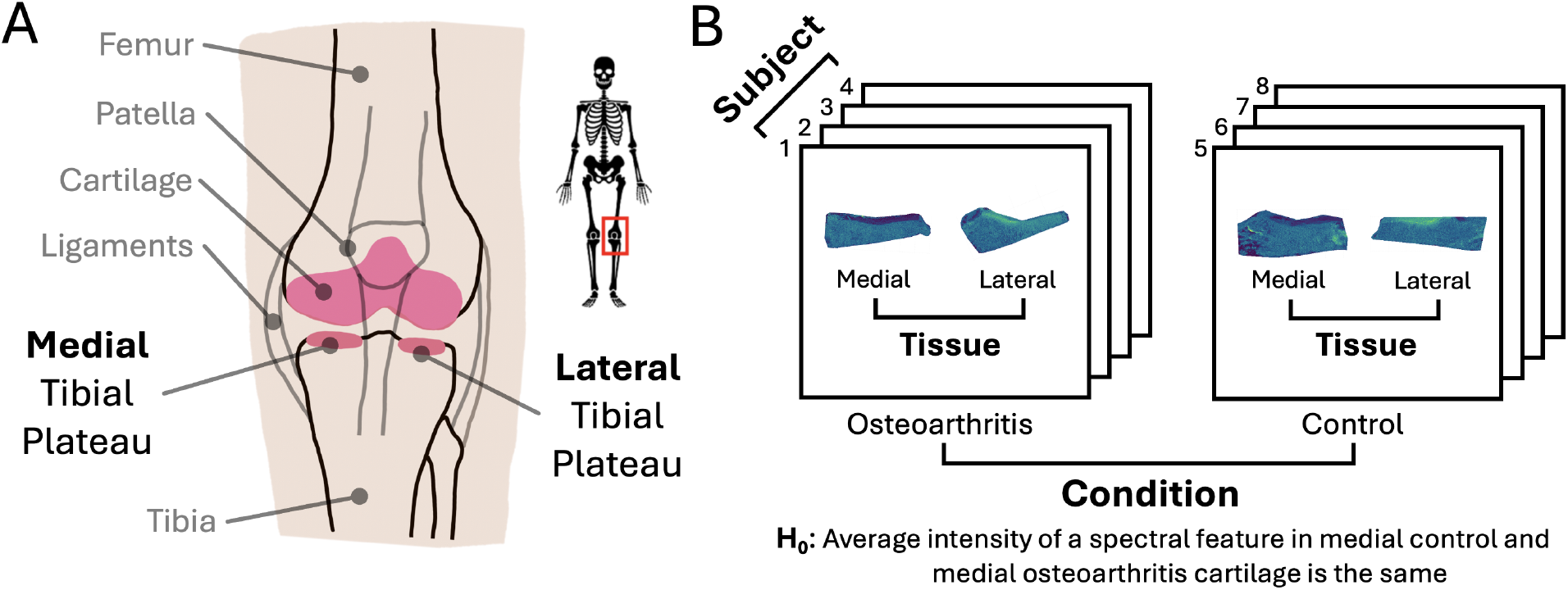
OA study: Biological context and research question informed the study design and data structures. **A:** Anatomy of the knee joint. **B:** Experimental design. Four subjects with and four without OA contributed lateral and medial tissues. One null hypothesis of interest is stated below.

Based on previously reported methods [16, 17], samples were fixed and demineralized before sectioning. Histologically-stained sections were optically imaged, revealing distinct cartilage pads above subchondral and trabecular bone. After paraffin removal and rehydration, samples were heated with citraconic acid for antigen retrieval. Samples were deglycosylated and digested with collagenase III before application of *α*-cyano-4-hydroxycinnamic acid imaging matrix and internal mass calibration standard [Glu1]-fibrinopeptide B[18] (reference m/z of 1570.676) of known concentration via automated sprayer. MSI was performed using a Bruker Scientific timsTOF fleX system, covering a mass range of 600–4000 m/z with a step size of 120 µm. Data acquisition yielded 16 raw imzML files, each with over 45,000 peaks, totaling to over 200 GB of data. Original data, as well as a reanalysis documented in this workflow, are publicly accessible via MassIVE (MSV000094448).

The statistical analysis workflow in this manuscript focuses on two main hypotheses. Hypothesis A assesses between-subjects differential abundance of analytes in medial tibial articular cartilage samples in two conditions, with versus without OA. Hypothesis B assesses within-subject differential abundance of analytes in medial versus lateral tibial articular cartilage samples with the OA condition. A secondary Hypothesis C investigates the synergistic interaction between tissue and condition. Complete methodological details of sample acquisition, data collection and analysis can be found in Schurman *et al*, 2026 [19].

### Simulated dataset 1

Simulated dataset 1 was designed to illustrate the impact of ROI segmentation on detecting differentially abundant analytes. Eight 100 × 100 peak picked images were simulated using the Cardinal package *Cardinal::simulateImage()*. The simulation mimicked peak picked images with 3000 centroided spectral features, Normally distributed peak and pixel intensity error, no multiplicative variance, moderate spatial autocorrelation, and no intensity-dependent missing values. For each image we defined a donut-shaped ROI comprising about one-third of the total area of the image. Pixels in the ROI of the first 100 spectral features had a mean difference from baseline of 40 to create a set of ROI-defining features shared by every image. The next 2400 spectral features of every image had no difference in mean intensity between conditions. To represent the differential abundance between conditions, the last 500 spectral features in images from Condition A had a mean intensity difference of 30 within the ROI. The simulation parameter values were based on the experimental dataset. The simulation code can be found in the vignette *Simulation-1*.

### Simulated dataset 2

Simulated dataset 2 was designed to illustrate the impact of biological and technical variation on differential analysis. Sixteen 100 × 100 images were simulated using the Cardinal package *Cardinal::simulateImage()* with an experimental design that mirrored the experimental dataset. The simulations mimicked peak picked images with 300 centroided spectral features, Normally distributed peak and pixel intensity error, no multiplicative variance, moderate spatial autocorrelation, and no intensity-dependent missingness. The simulation had approximately 4:1 average technical variation to average biological (between-subject) variation. Half of the spectral features were simulated to have no relationship between mean intensity, condition, and tissue. In the other half, mean intensity of the simulated features had a systematic, additive difference between conditions and tissues, and a random difference specific to each sample. The simulation was then repeated with the ratio of technical and average biological variation reversed to 1:4, such that the total variation remained approximately the same. The simulation parameter values were based on the experimental dataset. The simulation code can be found in the vignette *Simulation-2*.

### Overview of the proposed workflow

The proposed workflow for differential analysis of MSI experiments with complex designs is overviewed in Figure 2. It relies on the open-source R/Bioconductor Cardinal implementation, and is augmented by custom scripts involving other open-source R functionalities as necessary. Examples of the workflow in this manuscript were implemented using machines with Apple M-series processors and at least 64GB of memory, unless mentioned otherwise. The code is available at https://github.com/EBRogers/MSI-Arthritis-Vignettes and can be easily viewed at https://ebrogers.github.io/MSI-Arthritis-Vignettes.

**Figure 2:**
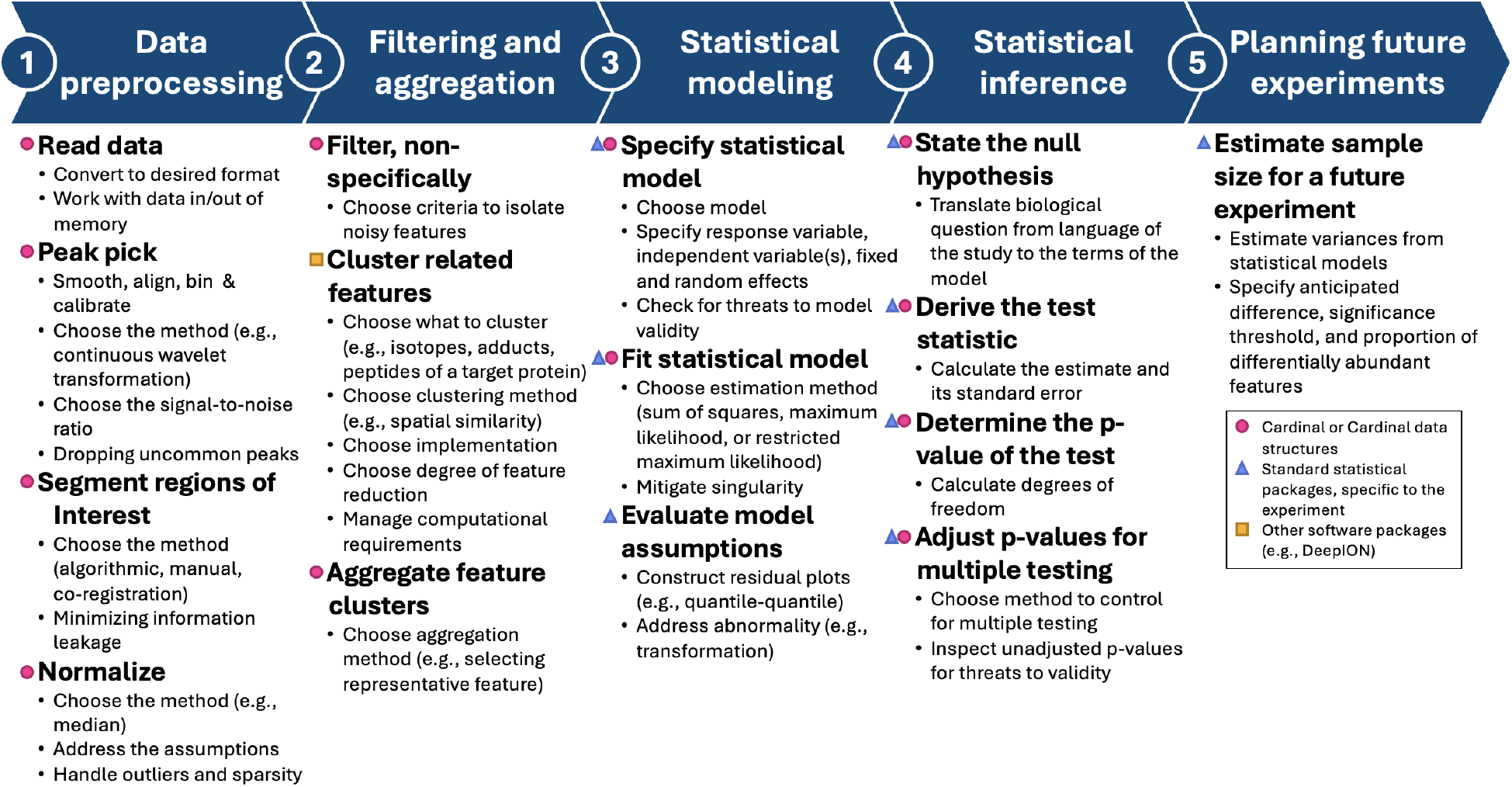
Proposed workflow for statistical design and differential analysis of MSI experiments with complex designs. Steps are labeled according to the availability of general open-source implementation in Cardinal, or other open-source implementations.

### Proposed workflow, Step 1: Data preprocessing

The first step, data preprocessing, aims to enhance the variation of interest in the mass spectra, and reduce the artifacts of the nuisance variation. This step is not specific to differential analysis and is common to most MSI analysis workflows. From the statistical perspective, it implements the quality control and data preparation that strongly impact the plausibility of the statistical modeling assumptions and of the outcome of differential analysis. Code for the proposed implementation of this step is in vignette *1-Preprocessing*.

#### Read Data

Cardinal takes as input arrays of processed or continuous MSI data. Cardinal supports.imzML [9, 20] files, a open source standard supported by most MSI software. Converters to .imzML from other formats are available online [9, 21]. Cardinal supports any ionization type or instrument vendor that can export data as or convert data to .imzML files, but is most frequently used for two-dimensional MSI data from MALDI and DESI sources. For example, the OA dataset was collected using a Bruker Scientific timsTOF fleX and files corresponding to the individual tissues were exported as separate .imzML files for input to Cardinal. *Cardinal::readMSIData()* loads the dataset. Internally, Cardinal uses the resulting data structures that store MSI data on the system hard drive to facilitate working with datasets larger than available memory.

#### Peak pick

Recalibration and alignment coerce peaks and features onto a shared mass axis between pixels and samples. *Cardinal::recalibrate()* aligns the mass spectrum of each pixel to a set of reference peaks, either m/z values of an internal standard or estimated using *Cardinal::estimateReferencePeaks()*. For example, with the OA dataset we smoothed, removed baseline, recalibrated the mass spectrum twice for each sample, first using a set of reference peaks estimated using the default parameters then again using the [Glu1]-fibrinopeptide B internal standard and four other endogenous reference peaks common to all samples. Each MSI in the OA dataset had around 45,000 pixels, making recalibration especially computationally intensive. Peak picking locates and quantifies peaks that likely belong to analytes, as opposed to noise, producing a filtered list subset of centroided spectral features as output. *Cardinal::peakProcess()* picks peaks and aligns centroid features using one of six supported picking methods: *diff, sd, mad, quantile*, and *filter*, which estimate noise using the derivative, standard deviation, mean absolute deviation, rolling quantiles, or dynamic filtering of local peaks respectively, as well as *cwt* that detects peaks with continuous wavelet transformation. For example, in the OA dataset we used a high signal-to-noise ratio of 7 and the diff peak picking method. The latter estimates noise based on the derivative of the signal, which captures local variations in spectral shape to differentiate true peaks from noise.

Spectral alignment after peak picking ensures that features of all spectra are aligned within each MSI. In the OA dataset, to ensure that all peaks were comparable between the samples, we combined samples into a single MSI where each sample was a run. We then performed peak picking and alignment with a high signal-to-noise (SNR) ratio of 7 to retain only the relevant peaks. By default, Cardinal picks on all pixels, but *Cardinal::sampleSize()* can also be used to pick on a random subset of peaks as a computational shortcut. For the OA dataset we opted to pick every pixel for thoroughness, we discarding any peaks found in less than 5% of spectra of any individual run. Preprocessing the 20 OA dataset, consisting of 16 samples and over half a million pixels took about 12 hours with parallel processing on a powerful laptop, with the recalibration steps being most time consuming.

#### Segment regions of interest (ROIs)

ROIs confine the analysis to more homogeneous sample regions. The ROIs can be constructed using information external to the mass spectra. For example, annotated histologically stained optical images, high-resolution spatial transcriptomics or other modalities can be coregistered with MSI using *MSIreg* [22], an open-source coregistration package compatible with Cardinal. The annotations can then be manually transferred from these modalities using *Cardinal::selectROI()* on any other visualization renderable by Cardinal’s visualization system. In clinical and translational studies of differential abundance, ROIs are ideally defined using coregistration of pathologist-annotated histology images. However, if external information is unavailable, an ROI can be obtained by univariate segmentation of one (or a small number of) representative MS features, such as known or putative ROI markers. The univariate segmentation can be achieved in Cardinal using spatial-Dirichlet Gaussian mixture models (sDGMM) with *Cardinal::spatialDGMM()* [23].

Many exploratory investigations use multivariate clustering techniques for MS image quality evaluation and discovery of biological structures. This is done using general-purpose methods such as K-means or UMAP[24], as well as methods specifically designed for MSI such as Spatial k-means[25] and Spatial Shrunken Centroids [26]. ROIs defined by these methods may represent latent biological structures of interest and can be further investigated by secondary re-evaluation by a pathologist, but they are unfortunately not fully compatible with downstream differential analyses.

Specifically, the differential analysis of ROIs defined by multivariate segmentation is compromised by two issues, namely double-dipping (also called data leakage) [27], and selection bias. These issues are illustrated in Figure 3 in a hypothetical (and exaggerated) example of intensity-based ROI in an MSI experiment with one spectral feature. Figure 3A illustrates double-dipping, which arises in experiments testing between ROIs in a single-sample, specifically when feature intensities are used twice: first to construct the ROIs of interest, and then to test the pixels between the ROIs for differential abundance. The segmentation produces ROIs with intensities that are separated by design, even if all the variation between the ROIs is due to random chance. The null hypothesis of no difference in average intensity between the ROIs is therefore not meaningful. The resulting p-values are invalid, tend to be too small, and produce false positive differentially abundant features.

**Figure 3:**
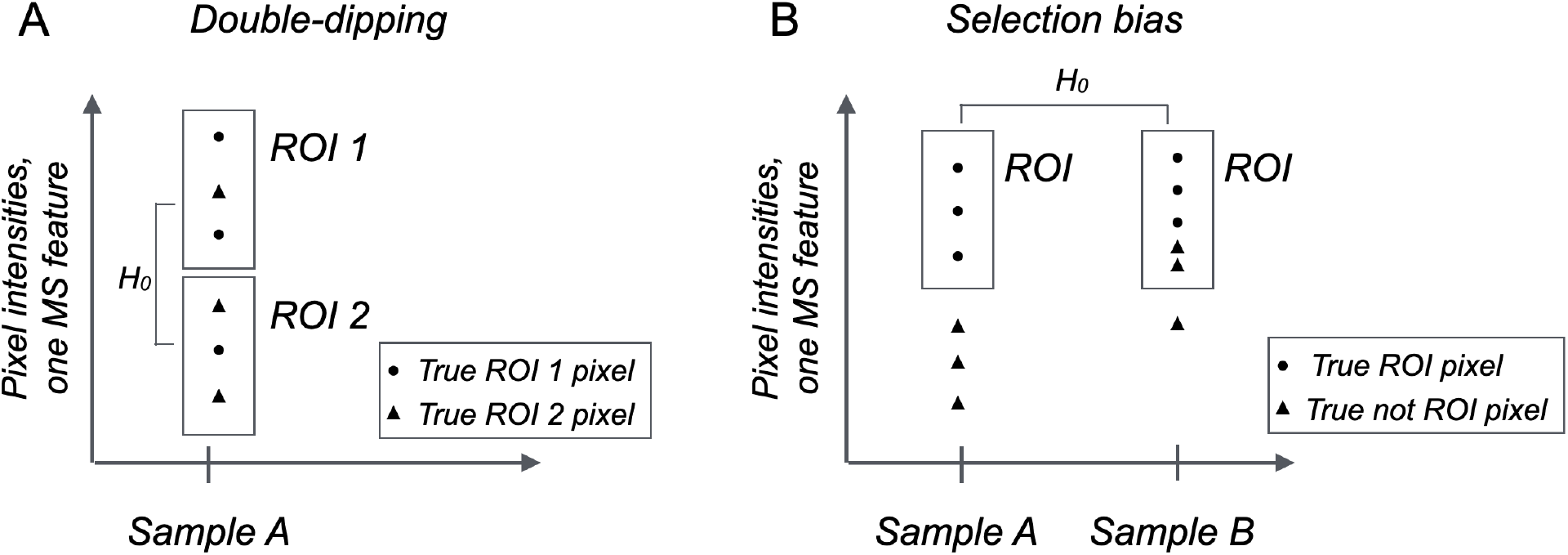
Hypothetical (and exaggerated) example of intensity-based ROI definition in an MSI experiment with one MS feature. Boxes denote ROIs selected based on intensity values. **A:** *Double-dipping. Intensity-based segmentation based on one feature in* one sample produces ROIs with intensities that differ by construction. Due to random variation in the intensities, the ROI may fail to correctly select the pixels. The null hypothesis of no differential abundance between RO1 1 and ROI 2 has no meaning and the resulting p-values are invalid. **B:** *Selection bias*. Intensity-based segmentation of two samples selects ROI with similar intensity ranges. However, due to random variation in the intensities, the ROI may fail to correctly select the pixels. This compromises the ability to detect differential abundance between the true ROIs.

Figure 3B illustrates selection bias, which arises in multi-sample experiments where the same feature intensities are used twice: first to independently select an ROI in each biological sample, and then test the selected ROIs for differential abundance between the samples. The segmentation produces intensity-dependent ROIs that, due to random variation, may fail to correctly select the relevant pixels. This in turn leads to ROIs with artificially similar average intensities and to false negative differentially abundant features.

The example in Figure 3 is intentionally extreme for the purposes of illustration. In experiments with a large number of features, the influence of the individual feature intensities is less dramatic, and the impact of double-dipping and of selection bias is more subtle and continuous. However, as we illustrate below, it has a non-negligible impact. Therefore, for the specific purpose of downstream differential analyses, we recommend selecting the ROIs based on external annotations, or when unavailable, using a small number of marker features that are subsequently excluded from differential analysis. Since selection bias is most relevant for multi-sample experiments with complex designs, we focus on this aspect in what follows.

The example OA dataset did not have histologically stained images for coregistration or annotations. Previous work has identified potential markers of cartilage [28]. One such marker, 1613.7666 m/z (a peptide of cartilage oligomeric matrix protein with sequence R.NALWHTGDTESQVR.L, matched to 1613.713 m/z in the OA study), demonstrated a spatial pattern consistent with cartilage in most samples, but was a poor candidate for univariate segmentation due to its low intensity. Instead, we selected the most colocalized ion with 1613.713 m/z (using *Cardinal::colocalized()*), 1141.545 m/z, for cartilage segmentation as it had high intensity and its spatial distribution consistently correlated with cartilage in all samples. We segmented the spatial distribution of this feature using Cardinal’s implementation of sDGMM [23] with adaptive weighting. Cartilage markers and subsequent ROIs were verified as representative of cartilage by subject matter experts. As a second step, the internal mass calibration standard [Glu1]-fibrinopeptide B (1570.679 m/z) spiked into the imaging matrix and six of its neighboring spatial analogs (1571.680 m/z, 1572.681 m/z, 1592.670 m/z, 1593.674 m/z, 1594.677 m/z, and 1608.659 m/z) were visually correlated to sample background areas, such as the intertrabecular space, negatively defining bone, potentially due to ion supression effects. We repeated the sDGMM-based single-ion segmentation on the pixel-wise mean of [Glu1]-fibrinopeptide B and its spatial analogs to further segment bone from background, limiting the segmentation to pixels not classified as cartilage. Separating background from tissue was necessary for later steps in the workflow such as filtering and aggregation. Finally, to avoid information leakage/confounding, we discarded the features used for segmentation from statistical analyses. Since the ROI segmentation only involved a few features, computational overhead was negligible.

#### Normalize

Normalization aims to eliminate technological artifacts between MSIs and pixels, and make them comparable. *Cardinal::normalize()* implements multiple normalization strategies, including *tic* for total ion concentration (TIC), *rms* for root mean square (RMS) normalization, or *reference* for reference normalization. Global normalization methods, such as TIC or RMS, can be performed on any dataset without addition of an internal normalization standard, but assume a constant baseline and a small number of differentially abundant analytes [29]. Reference normalization does not make any assumption regarding differential abundance but requires the presence of a reference standard with uniform concentration across all the pixels and samples [30].

Normalization can be undermined by sparsity of MSI measurements, i.e. by intensities of m/z features that are either missing or zero at some pixels. Sparsity arises when an analyte is absent in the pixel location, but may also be due to artifacts of data acquisition or processing (such as peak misalignment). Cardinal automatically stores MSI measurements as sparse matrices, filling missing values with zero. On one hand, excessive sparsity may bias summaries equalized by global normalization, such as the mean intensity in an ROI. On the other hand, excluding the sparse values from the normalization summary biases the summary towards large outliers. Normalization of MSI experiments, especially of experiments with multiple tissues, remains an active area of research [31]. In this manuscript, we recommend controlling sparsity when peak picking by using a high signal-to-noise ratio and excluding high sparsity features and pixels with few nonsparse peaks from analysis.

For example, in the OA dataset, all peak picked MSI had both high sparsity and many large outliers, compromising global normalization. Moreover, we observed both the high relative sparsity and pixel-to-pixel noise of internal mass reference standard within the cartilage ROI, which would degrade the quality of reference normalization with this standard. Therefore, we used a custom global normalization procedure, dropping sparse values and right-censoring pixel intensity above the 95^th^ percentile of each sample, prior to calculating the sample median as the normalization constant. The intensities were then scaled to the pooled median intensity of all samples. Pseudocode for this normalization procedure is available in Figure S2.

### Proposed workflow, Step 2: Filtering and aggregation

Feature filtering and aggregation reflect the statistical principles of defining the scope of comparison and performing quality control. They reduce the m/z features to a biologically or clinically relevant subset, and limit the multiplicity of hypothesis testing. Feature filtering removes uninformative features composed primarily of noise. Aggregation of redundant features, such as isotopes, adducts, and/or peptides of a target protein, reduces noise and facilitates detection of differential abundance. Since Cardinal does not have explicit functionality for this purpose, we propose a custom two-step process. We first eliminate non-informative features, and then cluster and aggregate isotopes to reduce redundancy and noise, as described below. The code for this step of the proposed workflow can be found in vignette *2-Filtering-Aggregation*.

#### Filter, non-specifically

Non-specific filtering eliminates uninformative m/z features, in a way that is blinded to the conditions of interest to differential analysis in order to avoid bias. For example, in the OA dataset we assumed that m/z with low intensity and variation contain limited biological information. *Cardinal::summarizeFeatures()* efficiently computes such summary statistics. We therefore calculated mean pixel intensity and standard deviation for each feature across all the samples, excluding the background and the missing values. We then removed features where both the mean and standard deviation fell below the 25^th^ percentile of all the m/z, as well as features with sparsity greater than 25% in any individual cartilage ROI. Such cutoffs are subjective, and their impact can be evaluated by sensitivity analysis.

#### Cluster related features

Clustering methods group m/z features according to shared characteristics such as spatial distribution, database-based annotation [32], mass error [33, 34], the Kendrick mass defect [35], or intensity ratios [36]. Most methods aim to cluster isotopes or adducts, however the type of similar features clustered depends on how clusters are identified. Cardinal currently does not offer a native feature clustering method, but offers several ways to export intensity matrices for input to external spatial clustering implementations [33, 37].

For example, in the OA dataset we clustered isotopes with similar spatial distributions using open-source software *DeepION* [33]. *DeepION* implements ResNet18-based encoders to generate denoised ion image representations. Then it traverses the mass scale, and evaluates theoretical mass differences and spatial similarity to identify possible isotopes. To adapt *DeepION* to multi-sample clustering, we combined all the images in a single data structure using Cardinal, while dropping background pixels to reduce memory usage. Unlike many other clustering methods, *DeepION* does not require users to pre-specify the number of clusters. However, it is computationally expensive. We used a 24-core cluster with 192GB of unified memory. Single-sample MSIs were processed in under 2 hours using GPU memory, while larger combined-sample images required several days.

#### Aggregate feature clusters

Aggregation combines the intensities of clustered m/z features into a single intensity value. Cardinal does not offer methods explicitly designed for feature aggregation. However, several Cardinal methods are easily combined for this purpose. For example, since not all isotopes and adducts are equally abundant, a simple aggregation represents a cluster with its most intense m/z feature. We implemented this approach in the OA dataset.

### Proposed workflow, Step 3: Statistical modeling

A fundamental statistical principle for studies of differential abundance is the explicit description of systematic and random sources of variation that affect the data, the hierarchy of these sources, and our assumptions. This description is accomplished using statistical models and is expressed in the language of probability. The models form the basis of statistical inference in the following Step 4, i.e., of the process that generalizes the conclusions beyond the subjects in the study to the broader populations of subjects. The models necessary for studies of differential abundance are distinct from models used for other goals such as class prediction or class discovery. Cardinal supports linear models appropriate for differential analysis using *Cardinal::meansTest()*, and exports data structures that are compatible with a wide range of other R-based implementations of statistical modeling. Code for this step can be found in vignette *3-Statistical-Modeling*.

#### Specify statistical model

In differential analysis of MSI experiments, different models are required for different experimental designs, and a separate model is typically specified for each m/z feature [38]. For example, in an experiment with two conditions, a linear model underlies the commonly used t-test. The model relates the variation in the dependent variable (the intensity of the m/z feature) to the variation in the independent variable (such as tissue or disease condition in the OA dataset). In statistical language, the variation between conditions is called “fixed effects”, reflecting the fact that the conditions are defined for the entire population of subjects, beyond the specific individuals included in the study. For example, in the OA dataset, Model 1 from Table 1 leads to a t-test comparing the average pixel intensities in an ROI between the control and OA conditions in medial tissues.

**Table 1:**
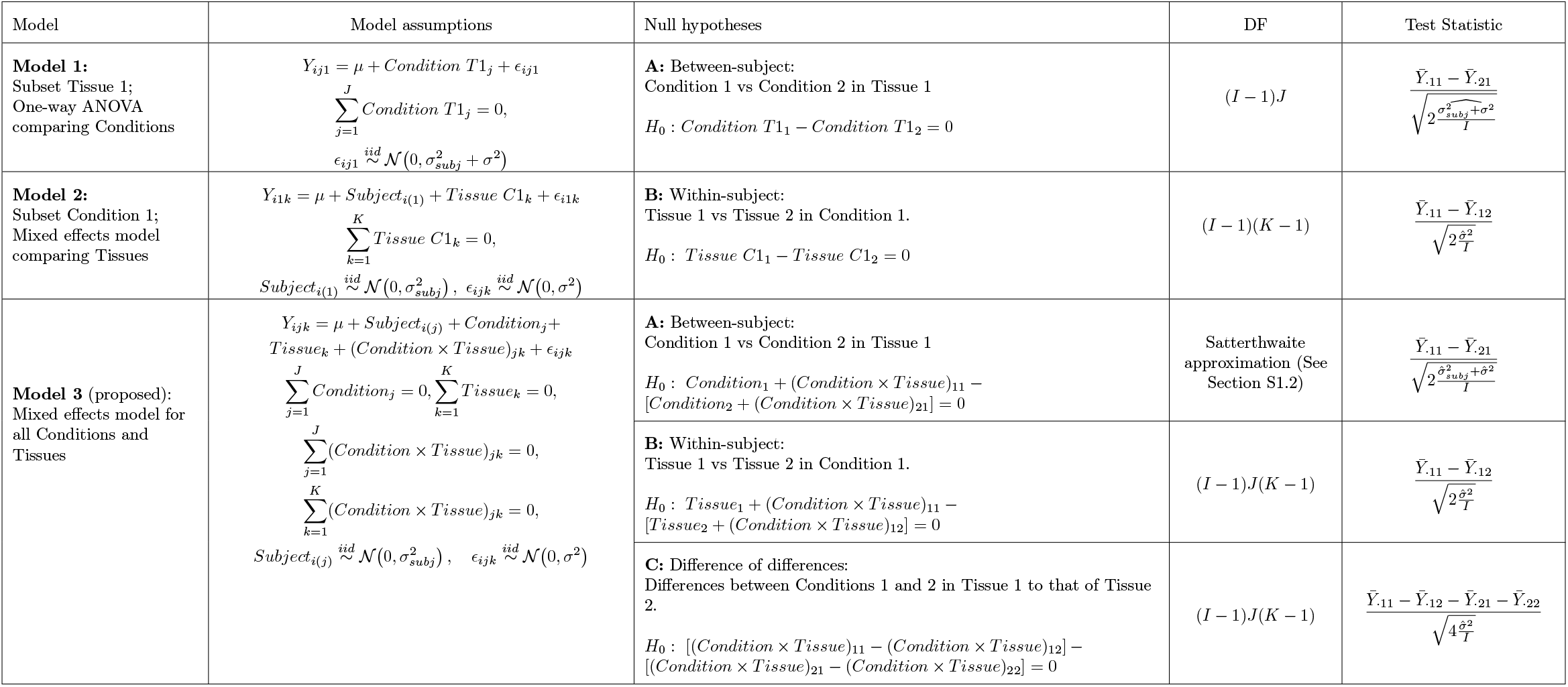
Statistical models for one m/z feature in the OA experiment. The models assume balanced designs and no missing values. *Y*_*ijk*_ denotes the mean intensity of the feature in the ROI of subject *i* = 1, *…, I*, condition *j* = 1, *…, J* (e.g, *j* = 1 for *control* and *j* = 2 for *OA*), and tissue *k* = 1, *…, K* (e.g., *k* = 1 for *medial* and *k* = 2 for *lateral*). Model terms with zero-sum constraints are fixed effects, and are the parameters of interest. Model terms that follow a Normal distribution are random effects. Model 3 is most comprehensive, and supports a wider range of null hypotheses. 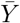 is the intensity averaged over the index indicated with a dot. 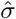 is the data-derived estimate of variation.

In experiments with more complex designs, linear models are readily extended to mixed-effects models. For example, in the OA dataset each subject has repeated measurements from lateral and medial tibial plateaus. Mixed-effects models distinguish the variation between subjects and the variation within repeated measurements on the same subject. Model terms reflecting subject-specific variation are called “random effects”, reflecting the fact that it is specific to the subjects randomly included in the study. For example, in the OA dataset, a mixed effects Model 2 from Table 1 leads to a paired t-test comparing the average pixel intensities in an ROI between medial and lateral tissues from a same subject with OA.

Linear models have also been extended with Empirical Bayes approaches, notably by *limma* [39], to capitalize on the large number of features and improve the characterization of random variation. However, the assumptions of *limma* are best compatible with relatively simple group comparison designs where each condition is represented by distinct sets of biological replicates, and are overly restrictive for experiments with paired and repeated measurements. Moreover, the benefits of the assumptions of *limma* diminish as biological replicates increase[40].

In MSI experiments, statistical models must also recognize the hierarchy of sources of variation affecting m/z features. Since intensities of m/z features are measured over a large number of pixels of a same sample, the pixel-to-pixel variation represents a lower level of hierarchy called subsampling (i.e., repeated sampling of a same biological replicate). It does not represent the biological between-subject variation, which is most relevant when comparing conditions. Mistaking the pixel-to-pixel variation for between-subject variation overestimates the evidence of biological replication, underestimates the extent of uncertainty, and produces overly optimistic results. For illustration, Table S2 in the supporting information contains examples of such models.

Finally, intensities of an m/z feature often correlate with the intensity in the neighboring pixels [41]. Models that explicitly account for both subsampling and correlation have been proposed [41, 42], however they are not easily scaled to complex designs or large datasets.

In this manuscript, we advocate for aggregating pixel intensities in an ROI, and reflecting the hierarchy of variation in these aggregates to accurately represent the experimental design, and to avoid scalability issues and restrictive assumptions. In the specific case of the OA experiment, we advocate for Model 3 in Table 1.

#### Fit statistical model

Since mixed-effects models are specified up to unknown parameters, the numeric parameter values are estimated from the data using maximum likelihood estimation (ML) or restricted maximum likelihood (REML)[43]. Cardinal automatically computes the mean intensity of a selected ROI, and fits statistical models with *Cardinal::meansTest(). Cardinal::meansTest()* automatically estimates model parameters with *stats::lm()* [44] for models with no random effects, or with either *nlme::lme()* [45] or *lme4::lmer()* [46] for models with random effects. Some m/z features may present edge cases, e.g. when the between-subject variation is negligible compared to the within-subject variation and measurement error, resulting in a singular fit [47, 48]. *Cardinal::meansTest()* handles this singularity by refitting a simplified model without the random effect.

#### Evaluate model assumptions

Statistical models describe the random variation in the data by postulating probability distributions. For example, Model 3 in Table 1 assumes subjects, *Subject*_*i*(*j*)_, are independent instances sampled from a Normal distributions. An important statistical principle is the evaluation of the plausibility of these assumptions based on the data, e.g. visually by means of diagnostic plots. Examples of such plots are shown in the Supporting Information Section S1.3. Some deviations from model assumptions can be addressed through transformations. For example, non-Normal distributions of the random terms can be addressed by log-transforming the intensities. The effectiveness of log transformation depends on the scale of the data and on the underlying processes driving skewness. In the OA study, log transformation did not improve the diagnostics, and was not pursued.

### Proposed workflow, Step 4: Statistical inference

Statistical models in Step 3 can be thought of as expert knowledge systems, and each comparison of interest can be thought of as a query. Statistical inference translates the scientific question into a null hypothesis expressed in terms of the model parameters. For example, in the OA dataset, the proposed Model 3 in Table 1 is queried with Hypothesis A for evidence of differential abundance between OA and control in medial samples. The same model is queried with Hypothesis B for evidence of differential abundance between medial and lateral tissues in OA. Models describing complex experimental designs support a wider range of queries. For example, in the OA dataset Model 3 in Table 1 supports Hypothesis C, which assesses the synergistic interaction (i.e., difference-in-differences) of feature abundances between tissues and conditions. Models that represent only one condition or one tissue type at a time (such as Model 2 and Model 3 in Table 1) do not support such a hypothesis directly.

Upon estimating the model parameters, statistical inference derives a model-based summary of the data called, test statistic, that is relevant to the null hypothesis. The test statistic typically takes the form of a signal-to-noise ratio such that its magnitude characterizes the extent of evidence against the null hypothesis. The test statistic is compared to a pre-defined cutoff to control the False Discovery rate, i.e. the expected proportion of false positives in the list of differentially abundant features. Fortunately, many statistical packages, including Cardinal, automate this process in a way that can be used by scientists with a limited statistical background. Code for this step in the special case of the OA experiment is in vignette *4-Statistical-Inference*.

#### State the null hypothesis

The null hypothesis compares the mean abundances of an m/z in the broad populations represented by the subjects in the study. It expresses these population-level means as a linear combination of the model parameters, called a contrast. For example, in the OA dataset, a query of Model 3 for evidence against null Hypothesis A (the difference between mean intensity of OA versus control medial samples) is written as

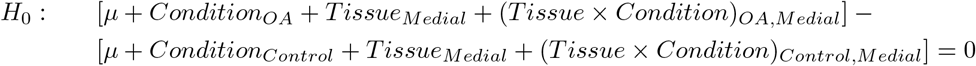

*Cardinal::contrastTest()* employs shortcuts to specify common comparisons of interest, and offers functionality to specify custom comparisons. We provide numerous examples of comparison specification in vignette *4-Statistical-Inference*.

#### Derive the test statistic

The test statistic is determined by the statistical model, by the parameter estimates, and by the null hypothesis. For example, in the OA dataset, in a special case of balanced design where every combination of conditions and tissues has the same number of biological replicates (e.g., for m/z feature with no missing values), a query of Model 3 for evidence against the null Hypothesis A is written in simple terms (Table 1). It is the difference of sample means 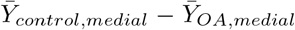 relative to the combined within- and between-subjects standard error of the group means 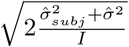. Similarly, a query of Model 3 against null Hypothesis B is the difference 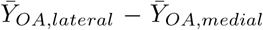 relative to 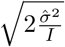. Note that Hypothesis B compares the lateral and the medial conditions represented by paired samples from the same subject, such that each subject serves as their own control. Therefore, Hypothesis B has a smaller denominator in the signal-to-noise ratio compared to Hypothesis A, reflecting a greater sensitivity of this comparison due to the nature of the design. The test statistics have a different form in unbalanced designs or in presence of missing values, and are automated in *Cardinal::contrastTest()*.

#### Determine the p-value of the test

To evaluate the extent of evidence against the null hypothesis, the test statistic is compared against a reference probability distribution. The reference distribution reflects the variation in the test statistic that we expect to see if the null hypothesis is, in fact, true. In linear mixed-effects models, such reference is the Student distribution, characterized by a parameter degrees of freedom. The p-value is the probability, calculated with respect to the reference, of observing a test statistic that exceeds in absolute value the test statistic that we actually observe. Small p-values indicate evidence against the null hypothesis.

The degrees of freedom quantify the amount of information available to test the null hypothesis, and larger degrees of freedom often give rise to smaller p-values. Calculating degrees of freedom is straightforward for many experimental designs and hypotheses (Table 1). Unfortunately, in some hypotheses, such as Hypothesis A from Model 3, degrees of freedom must be approximated, e.g. with the approach by Satterthwaite [49]. The approximation balances accuracy and computational efficiency, but can be undermined by numeric instability. (See Section S1.2 for more detail.) *Cardinal::contrastTest()* automates this step while accounting for the numeric edge cases.

#### Adjust p-values for multiple testing

Simultaneous hypothesis tests of many m/z features are bound to have at least some false positive rejections. Therefore, it is appropriate to control a multivariate false positive rate. A popular criterion is False Discovery Rate (FDR), i.e., the average proportion of false positive rejections of null hypotheses in a lists of differentially abundant analytes. The approach by Benjamini and Hochberg [50] transforms the observed p-values into adjusted p-values, such that the cutoff of the adjusted p-value is the FDR. *Cardinal::topFeatures()* transforms the p-values from hypothesis tests computed by *Cardinal::contrastTest()* according to the Benjamini-Hochberg approach.

### Proposed workflow, Step 5: Planning future experiments

Regardless of whether the current experiment produces significant findings, it contains useful information for optimizing the number of biological replicates in a future followup experiment [10]. Optimizing the number of replicates is important because under-replicated experiments lack sensitivity and overestimate the extent of the detected changes [51], while experiments with too many biological or technical replicates relative to biological and technical variation misallocate resources. In experiments where the number of replicates are fixed, such as when using tissues banked for research purposes, power calculations can inform experimenters of the minimum detectable difference, helping to contextualize results and inform resource allocation.

The optimal number of replicates depends on many factors. First, it depends on the statistical model and on the null hypothesis. Some hypotheses, such as Model 3 Hypothesis B in Table 1 in the OA dataset, use each subject as their own control. In this case, the test only needs to weigh signal against technical variation, instead of both technical and biological variation as would be the case if the hypothesis were comparing effects between different subjects. Because there is less noise to overcome, we anticipate that Hypothesis B requires fewer replicates than Hypothesis A to detect the same change in abundance. Second, it depends on operating characteristics, specifically on the smallest difference between the population means of analyte abundance Δ that we would like to detect, the largest FDR *q* and the largest probability of false negatives *β* that we are willing to tolerate, and the expected ratio between the number of analytes with no change in abundance to the number of differentially abundant analytes *m*_0_*/m*_1_. Finally, the number of replicates depends on the numerical estimates of the biological variation 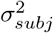 and technological variation *σ*^2^ expected in the future. These numerical estimates are derived from the current dataset, typically obtained as the median (or 75th percentile for a conservative estimate) over all m/z features. In situations where the number of replicates are fixed, these calculations can be used to estimate Δ instead.

#### Estimate sample size for a future experiment

In linear mixed effects models, the model and the null hypothesis define the standard error of a comparison. In a future experiment with one m/z feature the desired operating characteristics are achieved if the standard error is bounded by 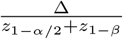 [10, 12, 13]. For example, in the OA dataset, Model 3 hypothesis A from Table 1 in the special case of a balanced design has the standard error 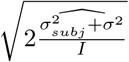, and the operating characteristics form the bound

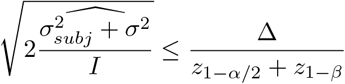

Solving for the number of biological replicates per condition results in

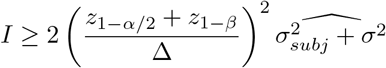

For experiments with multiple features we are interested in controlling the FDR. The original work by Benjamini and Hochberg[10] proposes to replace *α* with *α*_*ave*_

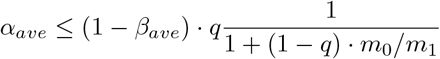

Although Cardinal does not offer native methods for sample size estimation, we implement these calculations as part of the proposed workflow. For example, in the OA dataset, we retained the median values of biological and technical variance across all the m/z features. We used these values to estimate the sample size for a range of percent differences and for different types of hypotheses. See vignette *5-Planning-Future-Experiments* for details of implementation.

## Results and Discussion

### In simulated dataset 1, Step 1 with all-feature ROI segmentation overfit the ROIs to noise

In the OA experiment raw MSIs had a noisy mean mass spectrum with over 45,000 peaks (Figure 4A). Compared to unprocessed images, peak picking as part of Step 1 significantly reduced the number of peaks, preferentially eliminating background and noisy peaks (Figure 4B). To evaluate the degree of noise reduction, we computed the coefficient of variation (CV) of pixel intensities from each feature before and after step 1. Preprocessing reduced the average pixel-wise CV across features from 3.54 to 1.81, indicating the preprocessed features had less pixel variation relative to their average intensity. Step 1 yielded a single image, with each sample a separate run, with a common m/z axis of 894 features and directly comparable intensities. Sections *Peak picking* and *Normalization* in vignette *1-Preprocessing* cover these steps.

**Figure 4:**
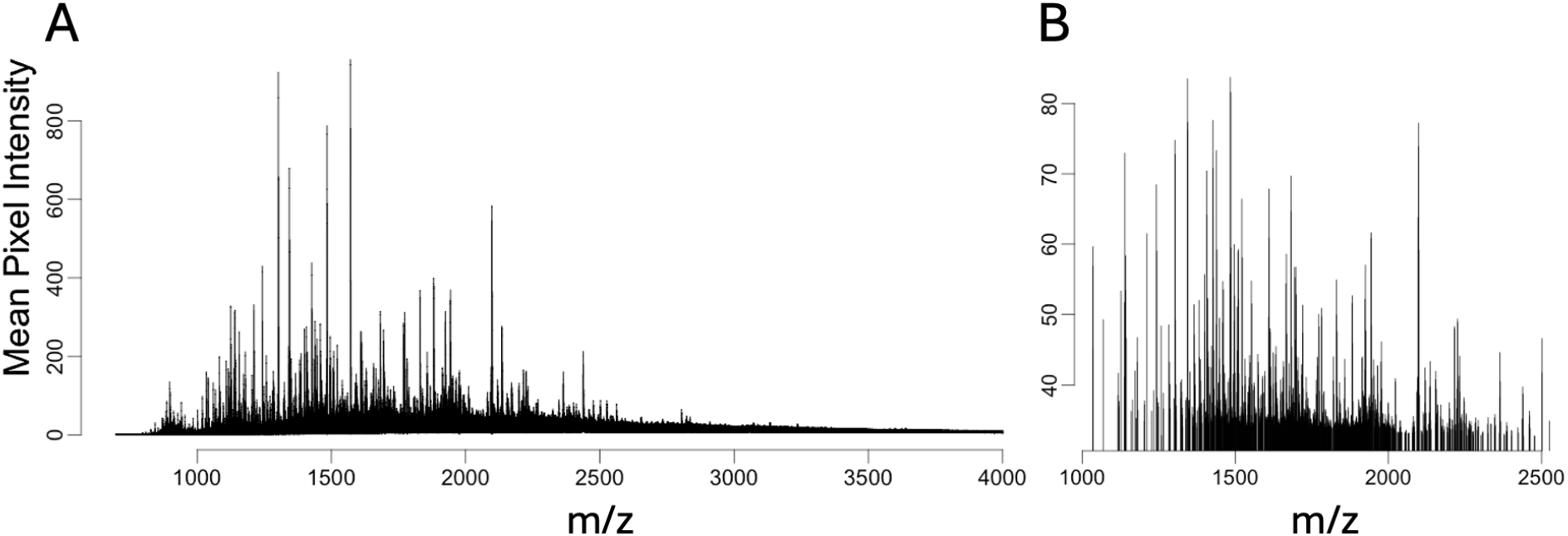
OA study: In Step 1, peak-picking and normalization removed background noise and reduced the overall number of spectral features. Note different scales of intensity axis, caused by scaling during normalization. **A:** Mean mass spectrum of sample 6891-OL before peak picking and normalization. **B:** Mean mass spectrum after peak-picking of peaks with high intensity relative to local noise and median normalization with Cardinal.

### In simulated dataset 1, Step 1 with all-feature ROI segmentation overfit the ROIs to noise

In absence of external information, multivariate segmentation is sometimes used to segment out ROIs capturing biological structures. We evaluated this approach using Simulated dataset 1 (Figure 5A) with known ground truth. We used the multivariate Spatial shrunken centroids (SSC) [26] and spatial K-means (SKM) [25] to separately segment each of the simulated images into 2 spatial regions.

**Figure 5:**
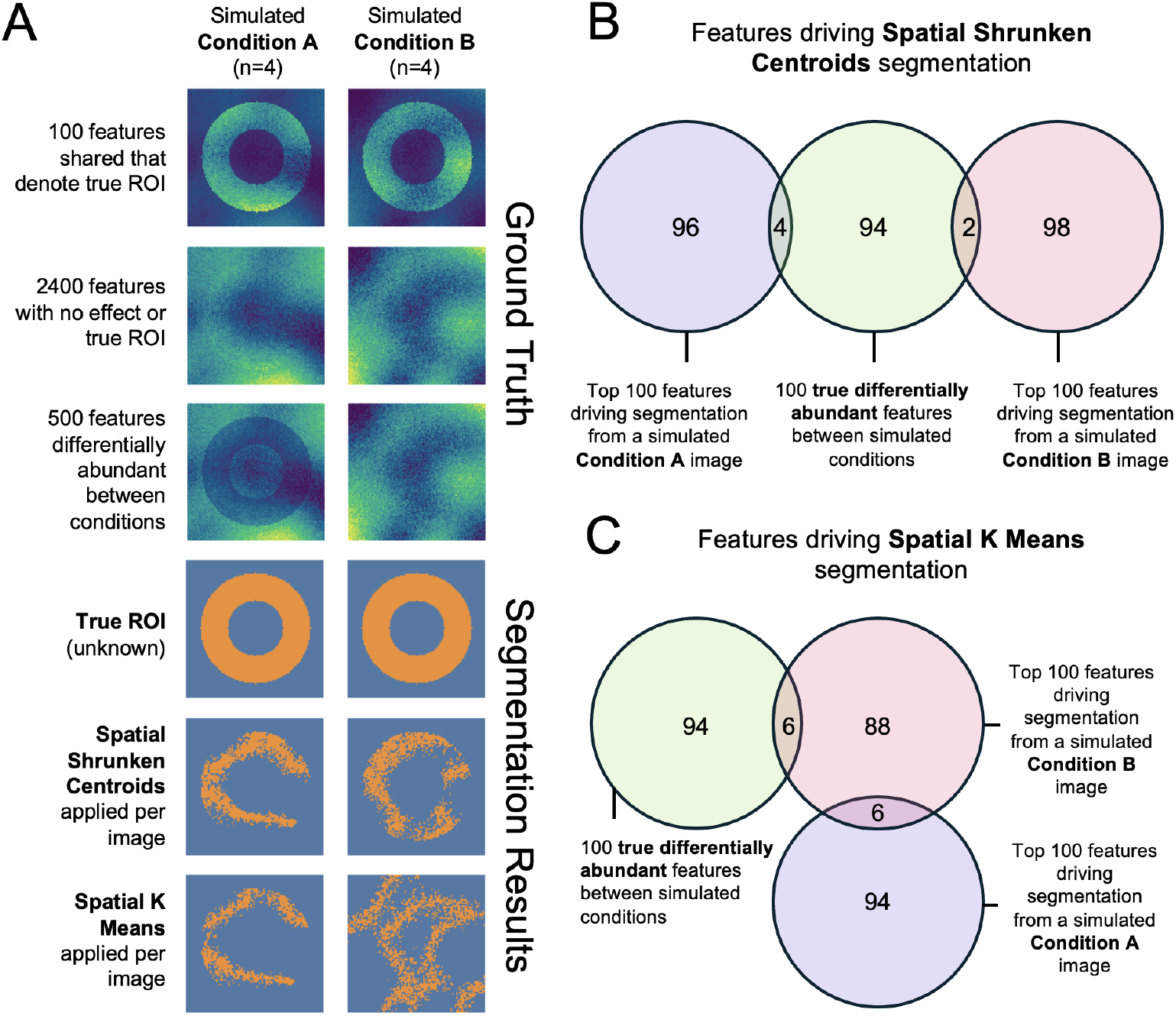
Simulated study: In Step 1, all-feature ROI segmentation with multivariate segmentation algorithms produced ROIs overfit to sample-specific noise. **A:** Simulation design and segmentation methods. All simulated images shared 100 donut-shaped ROI defining features, 2400 features with no differential abundance, and 500 features differentially expressed between simulated conditions only in the donut-shaped ROIs. The true ROI is shown, as well as the results from multivariate segmentation with SSC, or SKM. **B:** Venn diagrams of top 100 features defining the segmentation with Spatial Shrunken Centroids from a simulated condition A image, a simulated condition B image, and the 100 features that define true ROIs in both conditions. **C:** As in **B** but for SKM. Note that the none of the 100 ROI-defining features were in the top 100 features driving segmentation for the Condition A sample. All segmentation and visualization used Cardinal.

As seen in Figure 5B and C, SSC and SKM overfit to spatial noise, producing ROIs with homogeneous pixel intensities that inconsistently captured the true ROI. Although all samples shared the same 100 ROI-defining features, the features defining the segments differed between the methods and samples, and had little overlap with the truth (Figures 5B and C). Since the selected regions failed to capture the true ROI, and were composed of pixels with similar intensities, the t-tests for differential abundance between samples within these ROIs demonstrated substantially reduced the sensitivity (see Figure S1). Vignette *Simulation-1* offers code for this simulation.

### In OA dataset, ROIs from Step 1 highlighted the biologically plausible changes between tissues and conditions

As we did not have coregisterted images to use for segmentation, we defined ROIs in OA samples using nested single-ion (univariate) segmentation of representative markers of cartilage, bone, and background.

Features selected to segment ROIs exhibited representative spatial distributions of their tissue types and distinct distributions of pixel intensity after segmentation (Figure 6A). Single-ion segmentation produced intricate ROIs, defining the sponge-like structure of trabecular bone and cartilage pads (Figure 6B). By preselecting a limited number of features to define ROIs instead of segmenting ROIs using the entire feature set, we were able to obtain representative ROIs wile preserving features for testing of differential abundance later without the risk of confounding or information leakage. Cartilage and background ROIs segmented via the proposed method were highly similar to those generated via SCiLS in Schurman *et al* [19].

**Figure 6:**
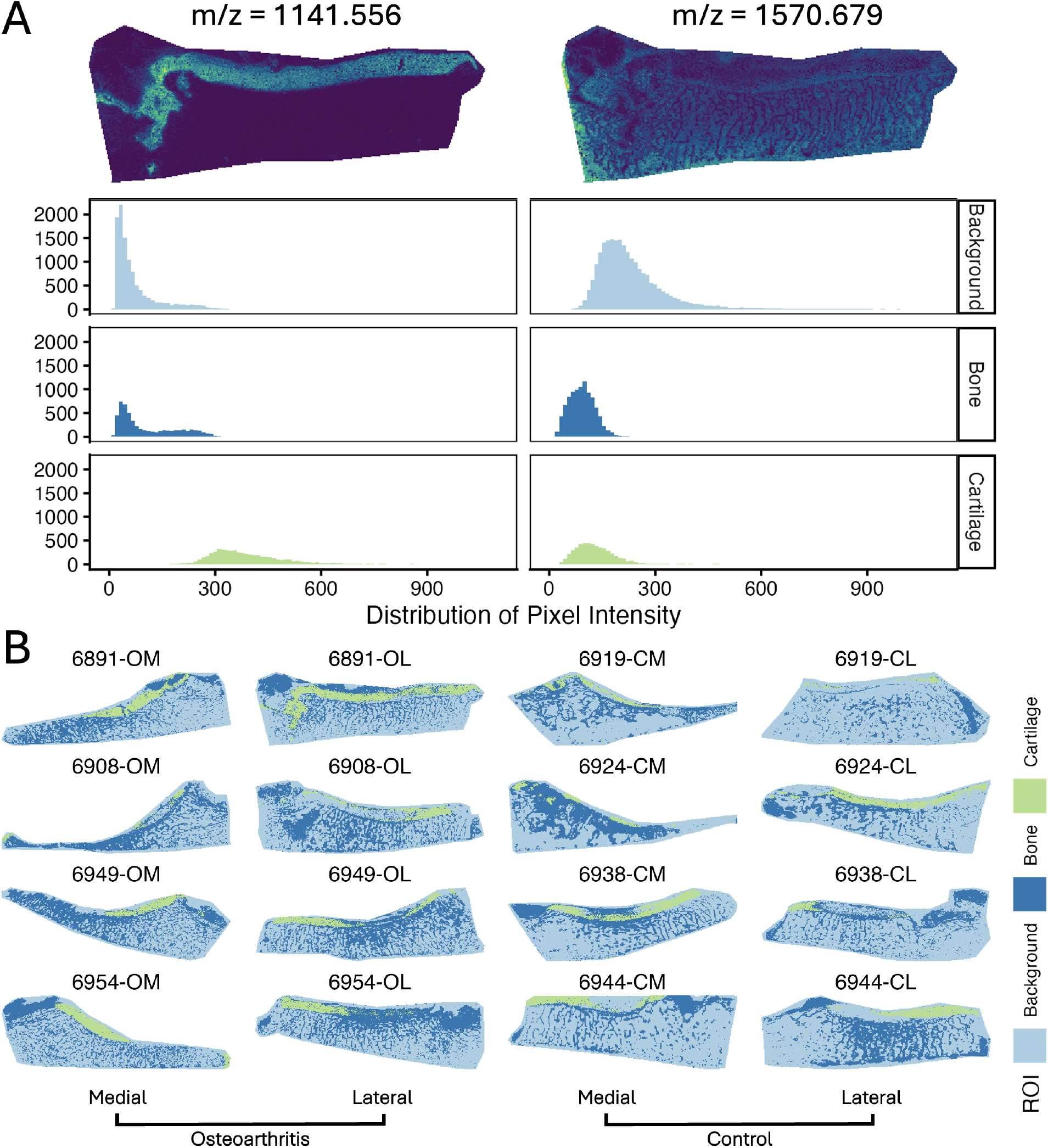
OA study: The choice of makers used to select ROI in Step 1 greatly influenced the ROI mean feature abundance. **A:** Single ion images and pixel intensity distributions of 1141.556 m/z and the internal standard ion 1570.679 m/z from sample 6891-OL. 1141.556 m/z accurately defined cartilage while the 1570.679 m/z internal standard added to the matrix distinguished tissue from non-tissue areas. **B:** The nested segmentation with sDGMM and ion markers highlighted the sponge-like structures of trabecular bone and the cartilage. Segmentation and visualization used Cardinal.

This nested segmentation required multiple iterations to identify the nesting order and the number of sDGMM clusters needed to preserve the structure of trabecular bone. This process consumed only a single feature, which was discarded from the subsequent analysis. The mass reference ions and related spatial analogs were also discarded. M/z values of the segmentation features can be found in Table S3. Section *ROI Segmentation* in vignette *1-Preprocessing* covers the proposed ROI segmentation procedure.

### In OA dataset, tests of differential abundance between conditions were more sensitive in cartilage ROIs defined with univariate segmentation of representative markers than multivariate segmentation

Algorithmically segmented ROIs can be defined using either univariate segmentation on a few representative features or multivariate segmentation across many or all features. We evaluated both segmentation methods for their effect sensitivity of statistical testing for differential abundance using the OA dataset (Figure 7). Cartilage ROIs were segmented with sDGMM univariate segmentation (described above) and again with multivariate SSC of the entire feature set. Mean intensity in cartilage was then tested between conditions of medial samples with t-tests.

**Figure 7:**
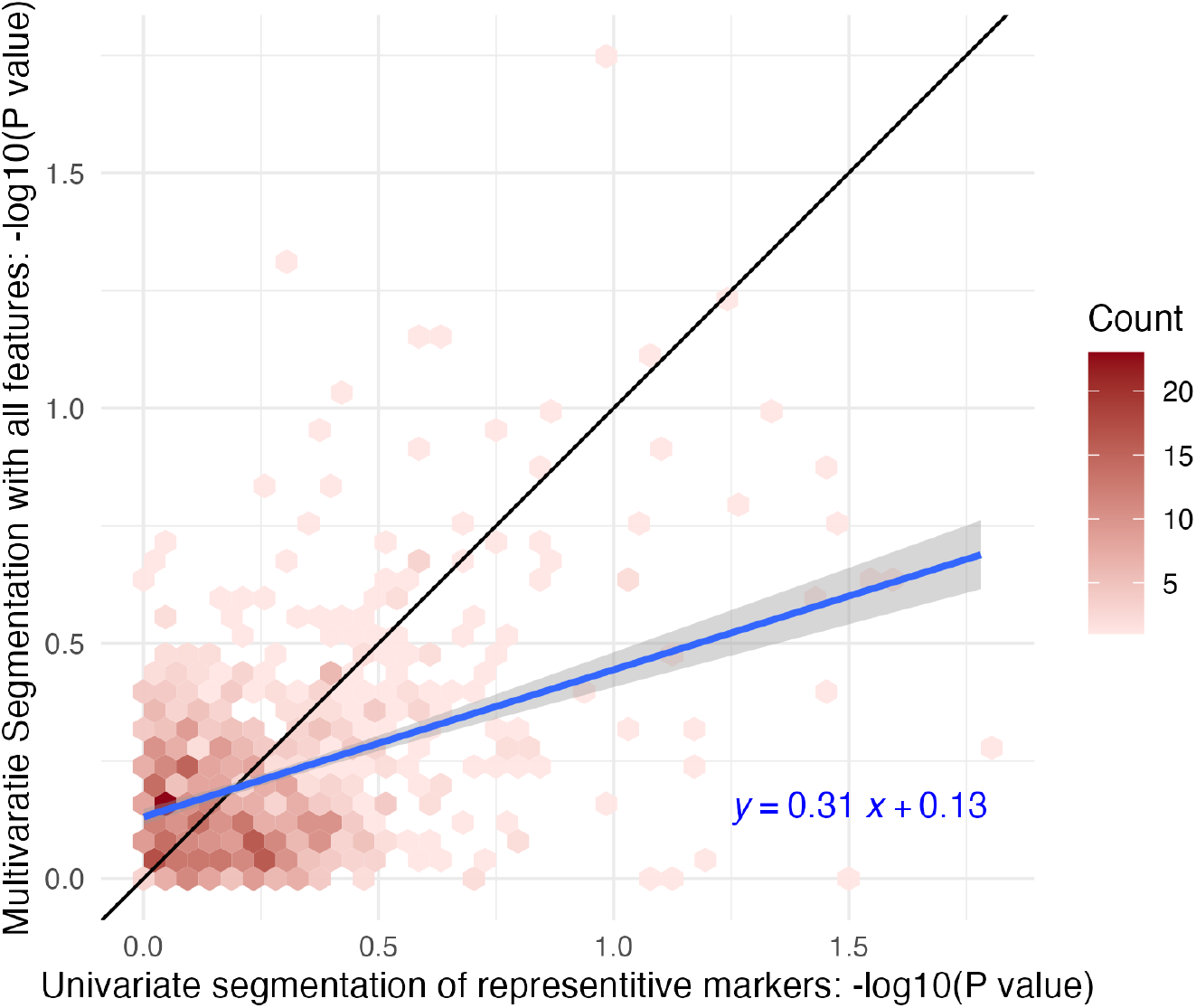
OA study: P-values from t-tests between conditions in cartilage ROIs defined by univariate and multivariate segmentation methods had substantial variation and, in multivariate segmentation, were on average more conservative. Two-sample t-tests of mean intensity in cartilage ROIs between conditions in medial samples were conducted for every feature. ROIs were defined by univariate segmentation of representative markers or multivariate segmentation of all features. The reference line in black indicates equal sensitivity. The fitted line in blue (with standard error) shows the correlation between p-values from each segmentation method. Tests were conducted on features before implementation of Step 2 and were not corrected for multiple testing.

As seen in Figure 7, tests conducted between cartilage ROIs created by multivariate segmentation were, on average, more conservative than tests between ROIs segmented with sDGMM, as evidenced by larger p-values. This effect was potentially due to the selection bias introduced when the same features used for segmentation were also used for statistical testing, as occurred with SSC (see Figure 3B). Both methods demonstrated high variation, and the conservative effect of multivariate segmentation was modest, highlighting the many inputs affecting sensitivity and algorithm performance. Vignette *1-Preprocessing* offers code for this simulation.

### In OA dataset, Step 2 reduced the number of hypothesis tests

After peak picking, discontinuation of analysis of the features used for ROI segmentation, and non-specific filtering, 684 features remained in the OA dataset. (See Figure 8A for contribution of each step.) Clustering of isotopes with *DeepION* and aggregation yielded 582 total features comprised of single ion features and representative feature aggregates. (See Table S4 for specific features clustered by *DeepION* and Figures 8B and C for examples of isotope pairs and triplets.) Finally, features with more than 25% sparsity in the cartilage ROI of any individual sample were discarded, leaving 124 features for further analysis. (See Table S5 for features dropped during sparsity filtering.)

**Figure 8:**
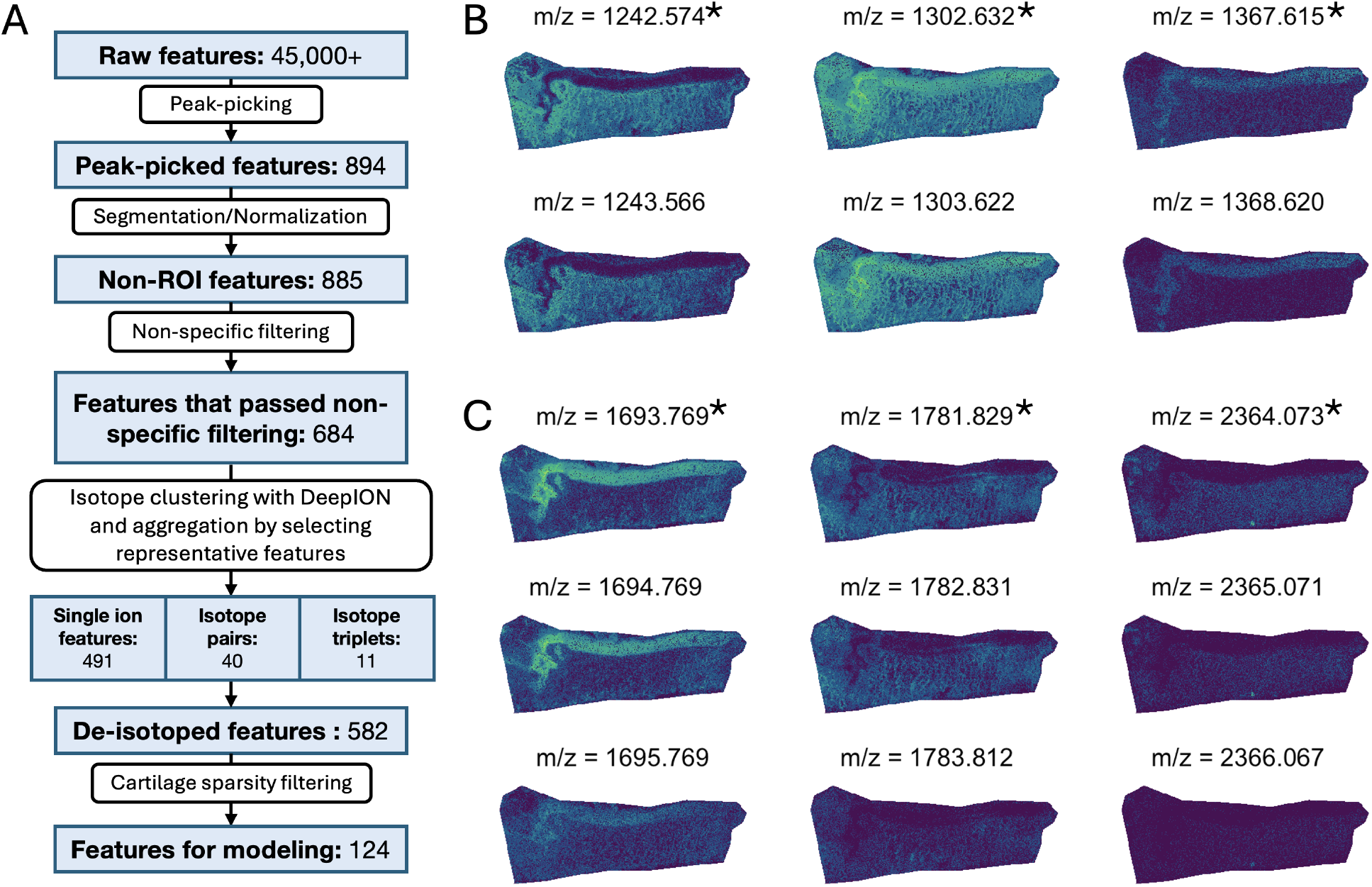
OA study: Step 2 reduced the number of features without obscuring their relevance. **A:** Workflow of reduction in features from peak picking, segmentation/normalization, non-specific filtering, feature clustering and aggregation, and cartilage sparsity filtering. Peak-picking occurred in *Cardinal* and clustering was done with *DeepION*. **B:** Isotope groups with two members in tissue 6891-OL had similar spatial distributions. Starred m/z are chosen as the representative of the isotope group. **C:** As in **B**, but for isotope groups with three members.

In the OA dataset, we filtered on both low standard deviation and low mean intensity preferentially filtered out features with low spatial information. Additionally, stringent sparsity-based filtering reduced the number of models with poor fit and minimized the bias towards outliers introduced by dropping sparse values at the time of normalization.

Even with modifications to decrease memory overhead *DeepION* required a large amount of memory and computation time, especially when run without GPU support. Additionally, since *DeepION* only has a prototype implementation, its application to new experiments may require computational skills. As ease of use remains a limiting factor for all open-source software, additional developments are needed to streamline the analyses. The filtering and aggregation steps are presented in vignette *2-Filtering-Aggregation*.

### In OA dataset, diagnostics in Step 3 protected against reporting unreliable results

To assess differential abundance in the OA study, we modeled mean intensity of features with Model 3 from Table 1. For about a third of features modeled, (41 of 124), subject variance was too small to accurately distinguish the between-subject variance from residual variance. This phenomenon is known as model singularity. We used *lme4::lmer()*’s default remediation strategy, refitting the model without the random effect, as the single-term random effect of subject in Table 1 was already the simplest structure. Singular models with more complex random effects should be refit with more parsimonious effects structures.

All 124 models were assessed for Normality of residual errors, similarity of variances between conditions and tissues, and presence of residual outliers. We retained only models satisfying these assumptions, reducing the chance of reporting false positive or negative results and the number of statistical tests requiring correction. An example of a model satisfying residual assumptions is showcased in Figures 9A through D. (Examples of poor fit are showcased in Figure S3.) The deviations of model assumptions were most frequently associated with highly sparse data. Thus the diagnostic evaluation of the model assumptions provided insights into threats to validity and reasons for potential lack of sensitivity and unreproducible results. (See Table S6 for evaluations of model assumptions and specific features retained for statistical inference.) These steps are covered in vignette *3-Statistical-Modeling*.

**Figure 9:**
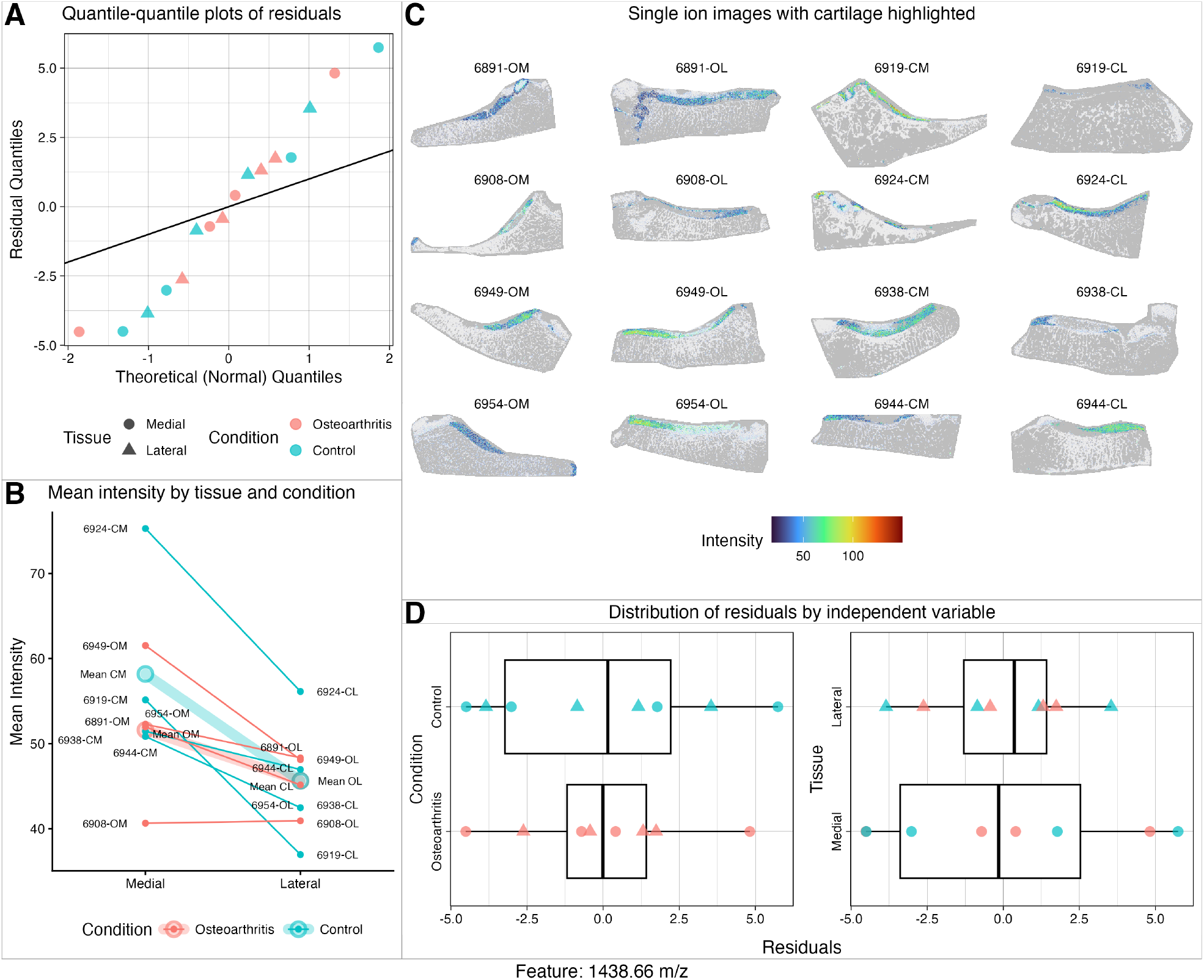
OA study: The model fit to 1438.66 m/z in Step 3 satisfied the assumptions of residual Normality, equal variance, and lack of significant outliers. **A:** Quantile-quantile plot is linear. **B:** Interaction plot visualizing the effect of tissue and condition on abundance by subject. **C:** Single ion images of each sample. Note low sparsity in cartilage ROIs. **D:** Box plots of residuals by tissue and condition.

### In simulated dataset 2, within-subject comparisons made in Step 4 were more sensitive detecting differentially abundant features

To demonstrate the impact of sources of variation in the data on the sensitivity of comparisons of interest, we simulated 16 MSIs with low biological and high technical variation, and the same 16 MSIs with the reverse, using the procedure detailed in the methods for Simulation 2. We fit each model specification from Table 1 to Simulated Dataset 2 using the methods outlined in Step 3 and assessed the sensitivity of Hypotheses A and B. We performed this procedure independently for each variance condition. We did not perform multiple testing corrections for this simulation. As Models 1 and 2 are equivalent to two-sample and paired t-tests respectively, this analysis essentially compared the sensitivity of t-tests investigating a single between- or within-subjects comparison to the same comparison from a mixed effects model using all available data, under different predominating sources of variation.

Interaction plots in Figure 10A illustrate the impact of variation source on mean intensity: either greater technical variation (*left*) or greater biological variation (*right*). When biological variation was high relative to technical variation (right), within-subjects comparisons from models accounting for covariance between subjects had greater sensitivity than between-subjects comparisons. (Figure 10B) Additionally, across both variance conditions, mixed effects models (Table 1 model 3) had greater sensitivity than models equivalent to paired and two-sample t-tests (Table 1 Models 1 and 2, respectively) regardless of dominant variation source. This was due to mixed effects models’ use of both independent variables and all observations, increasing the degrees of freedom and precision of variance estimates. Section *Dominant variance and sensitivity* in vignette *Simulation-2* offers code to reproduce these analyses.

**Figure 10:**
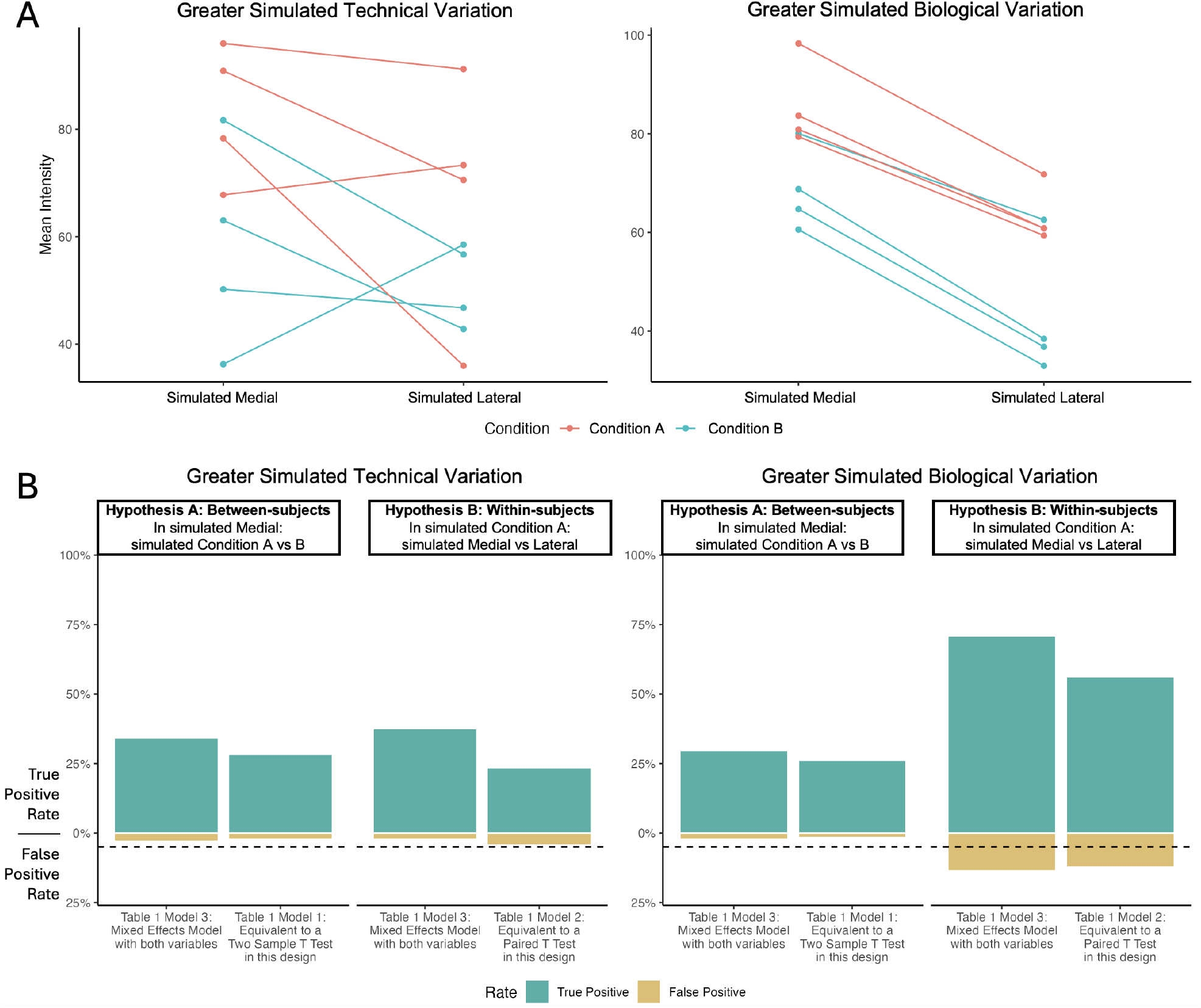
Simulated study: In Step 4, detecting differentially abundant features was more sensitive in within-subject comparisons (medial vs lateral) in than in between-subject comparisons (condition A vs condition B) in presence of large biological variation. **A:** Interaction plots of two representative simulated spectral features. *Left:* within-subject and technical variation larger than the biological between-subject variation. *Right:* within-subject and technical variation smaller than the biological between-subject variation. **B:** True and false positive rates for 300 features simulated as described in Section, analyzed with models in Table 1. Within-subject comparisons of medial versus lateral tissues were more sensitive then between-subject comparisons of osteoarthritis versus controls in presence of large between-subject variation, since each subject served as its own control. Mixed effects models representing all conditions and tissues were more sensitive than models based on subsets of data in presence of high technical variation. Horizontal dashed line represents a false positive rate of 5%. Comparisons were considered positive when unadjusted *P <* 0.05.

### In simulated dataset 2, using pixels as replicates in Steps 3 and 4 produced many false positive differentially abundant features

To illustrate the impact of using pixels as biological replicates, we repeated the analysis presented above using models from Table S2 that specify using pixels as replicates. Although in the previous analysis Model 2 from Table 1 investigating Hypothesis B was equivalent to a paired t-test, Model 2 from Table S2 is equivalent to a two-sample t-test as pixels locations cannot be paired between different samples.

Models 1 and 2 from Table S2 had significantly greater rates of false and true positives than their counterparts that did not use pixels as replicates, regardless of the comparison or dominant source of variation. (See Figure 11.) Mixed-effects models using pixels as replicates for within-subjects comparisons also had high rates of true and false positives. In general for most comparisons and models, using pixels as replicates increased sensitivity such that any observed difference was significant. Between-subjects comparisons from model 3 from Table S2 showed high rates of true positives and low rates of false positives because subject-level variation was distributed across both subjects and pixels. Therefore increasing replicates (pixels) did not decrease the contribution of subject-level variance to the standard error. See Section S1.2 for details on the Satterthwaite approximation for degrees of freedom. Section *Pixels as replicates* in vignette *Simulation-2* offers code to reproduce these analyses.

**Figure 11:**
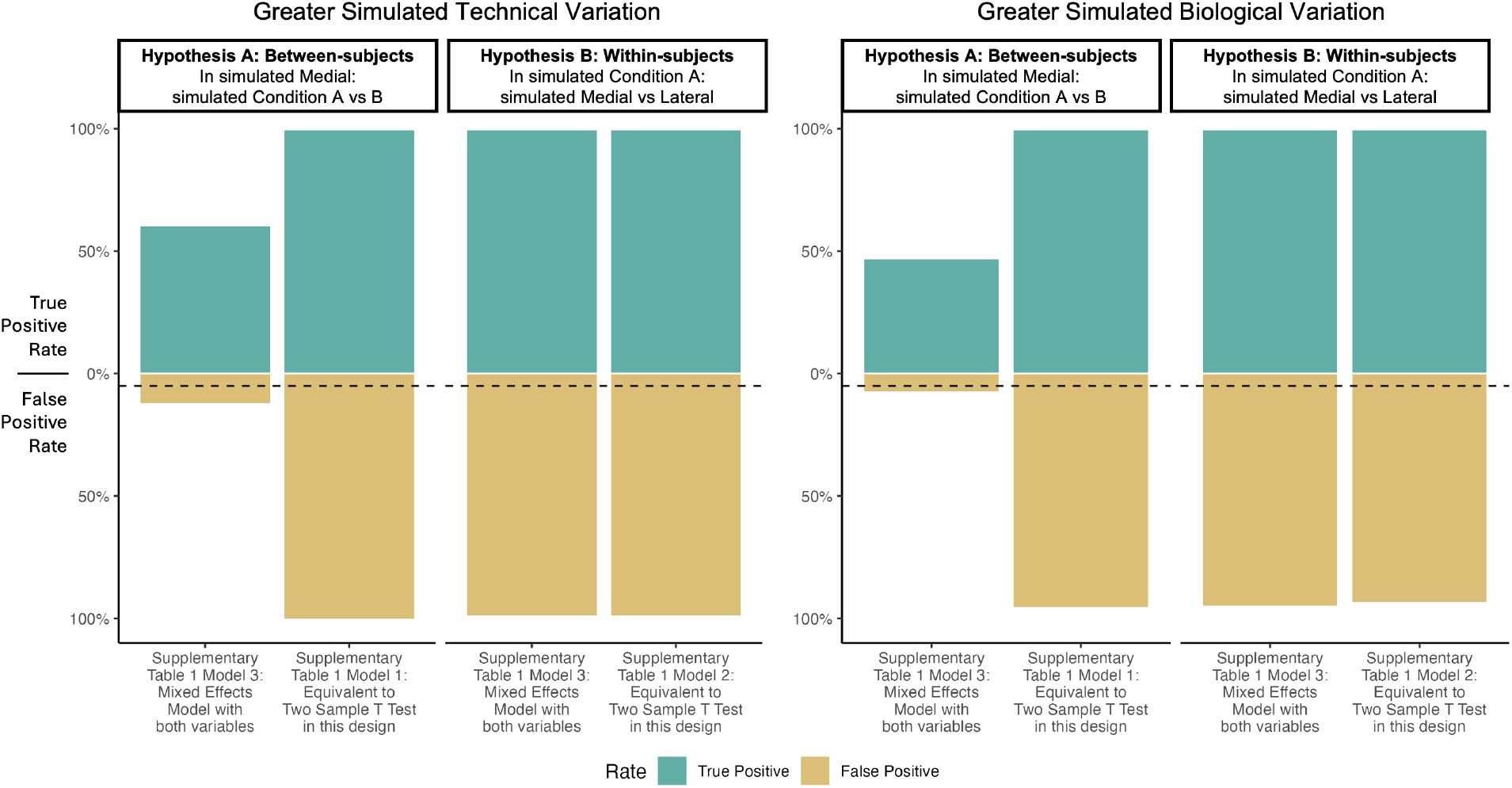
Simulated study: Using pixels as biological replicates in Steps 3 overfitted the data and produced many false positive differentially abundant features in Step 4. True and false positive rates for 300 features simulated as described in Section, analyzed with models from Supplementary Table S2. *Left:* within-subject and technical variation larger than the biological between subject variation. *Right:* within-subject and technical variation smaller than the biological between subject variation. When compared to Figure 10B, modeling feature intensities in each pixel, and viewing features as replicates, had a much higher rates of false and true positives than models using mean pixel intensity. Horizontal dashed line represents a false positive rate of 5%. Comparisons were considered positive when unadjusted *P <* 0.05.

### In OA dataset, Step 4 did not detect any differentially abundant features in cartilage

After multiple testing correction, there were no features with enough differential abundance to substantially overcome noise for either hypothesis. (See Figure 12A.) Thus, if differences do exist, larger investigations with more replicates are needed to identify them. We report estimates, standard errors, and unadjusted and FDR-adjusted p-values of Hypotheses A and B for Model 3 from Table 1 in Tables S7 and S8 respectively.

**Figure 12:**
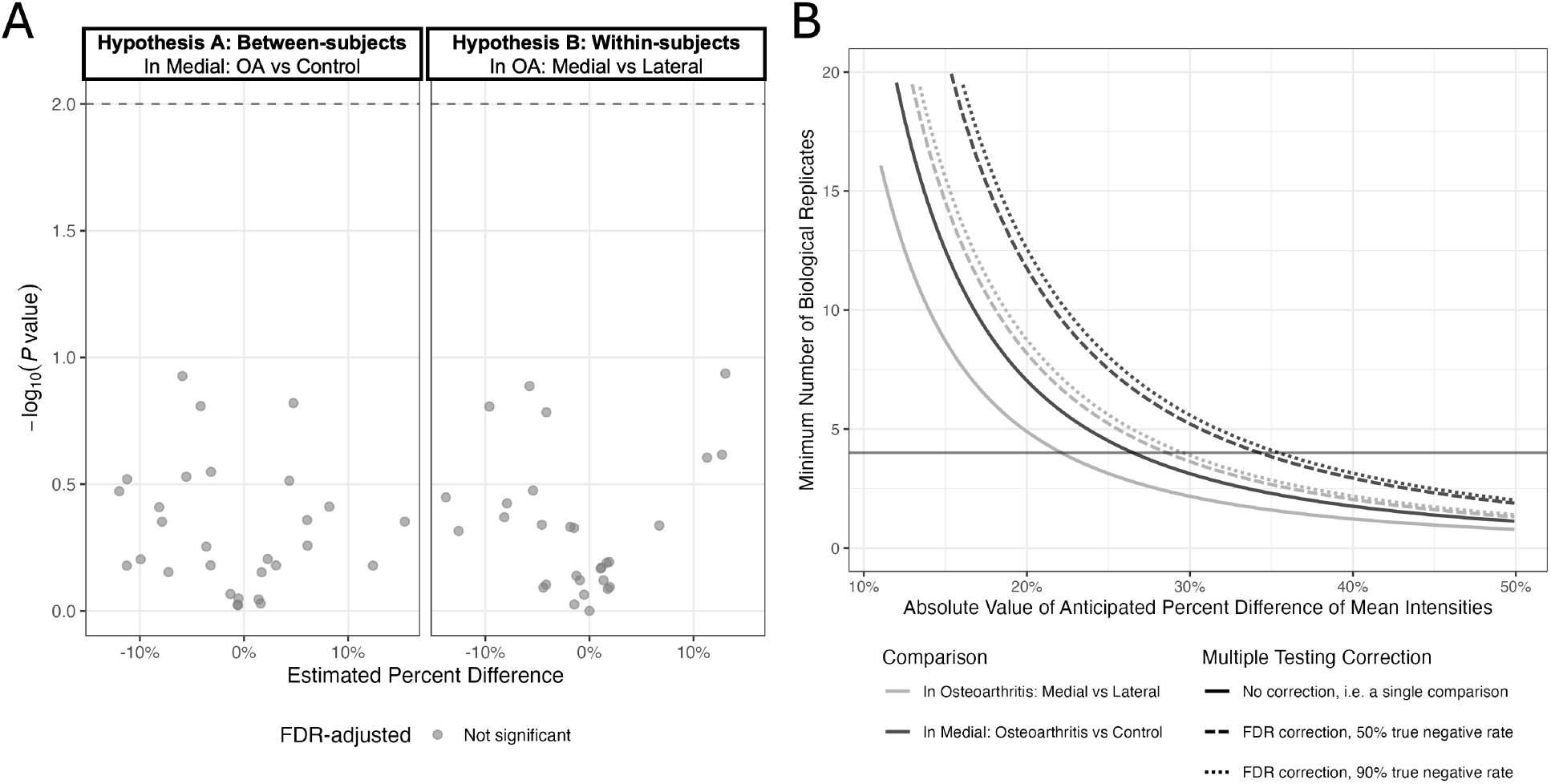
OA study: In step 4 and 5, within-subjects comparisons required fewer biological replicates to detect the same effect than between-subjects comparisons. **A:** Step 4: Volcano plots for 29 features. Gray line delineates unadjusted *P* = 0.01. No features remained significant after multiple testing correction at FDR-adjusted *P <* 0.10. **B:** Step 5: Minimum number of biological replicates needed to detect comparison percent difference based on the OA study dataset. The solid black line marks the number of biological replicates in the OA study, 4. Sample size calculations assumed power of 0.9 and *α* of 0.05 before correction.

In conclusion, multiple testing correction is sensitive to the total number of tests, and a larger number of analytes leads to more conservative thresholds. Eliminating the redundant analytes is therefore beneficial from the multiple hypothesis testing perspective.

### In OA dataset, Step 5 yielded estimates of sample size for future experiments

In the OA study, the minimum detectable difference was ±30% for within-subjects comparisons and ±36% for between-subjects comparisons, assuming median variance and 10% true positive rate. Figure 12B details the minimum number of biological replicates needed to detect a specified difference in the OA study experimental design for no correction and FDR-based multiple testing correction, assuming either 50% or 90% of features were not differentially abundant.

Increasing the assumed number of significant comparisons to 50% (dashed lines), increasing the number of biological replicates, or erroneously not correcting for multiple testing (solid lines) lowered the minimum detectable effect for both comparison types. As predicted, we observed increased sensitivity in within-subjects comparisons. These steps are covered in vignette *5-Designing-Future-Experiments*.

In conclusion, estimated sample sizes are most generalizable when preprocessing steps (Step 1) are replicated, as these steps directly influence estimates of variance.

### Summary of recommendations

#### Step 1: Data preprocessing

In addition to *randomization* and *replication*, preprocessing is essential to reduce noise and bias in MSI datasets. We recommend smoothing, baseline reduction, and recalibration of samples individually to reduce computational overhead, performing recalibration twice with both estimated peaks to improve pixel-to-pixel mass calibration and internal standards to import sample-to-sample comparability. Samples should be combined, picked, and aligned together to ensure a shared feature set. Ideally the entire sample area is picked, but a random subset of pixels can be used for homogeneous samples. For experiments investigating differential abundance, we recommend using exogenous information to segment ROIs and avoiding multivariate segmentation algorithms that use all or many features. We caution against using the same features for ROI segmentation and testing for differential abundance to avoid double-dipping or selection bias. We recommend choosing a normalization method with assumptions that best fit the experimental data. For experiments expecting few differentially abundant features, we recommend using median normalization as the median is less sensitive to outliers and sparsity.

#### Step 2: Filtering and aggregation

We recommend using non-specific filtering to drop noisy features based on conservative criteria with a clearly articulated set of assumptions and blinded to experimental variables. We also recommend applying aggressive sparsity filters to minimize the impact of sparsity on modeling on the *distributional assumptions* of downstream modeling. We recommend choosing the clustering method based on the experimental procedure, kinds of tissue in the sample, and expected types of related features in the sample. Methods that cluster all samples simultaneously are preferred. We advocate for aggregating clusters of similar features by selection of representative features for isotopes and adducts and pixel-wise mean for large feature groups.

#### Step 3: Statistical modeling

Choices such as model specification and type of model should be driven by the research question, hypotheses, and assumed structure of the data. We recommend modeling features independently using linear models that use all observations and specify all independent variables and subject as a random effect. We discourage use of pixels as replicates and advocate for use of mean pixel intensity in each ROI. We recommend using residual quantile-quantile plots and residual distributions by independent variable to evaluate model *assumptions* and applying remedial measures such as log transformations or weighted regression to address serious deviations from Normality. Further inference should be avoided on models that seriously deviate from assumptions.

#### Step 4: Statistical inference

We recommend testing of a limited number of hypotheses using signal-to-noise ratios (analogous to t-tests) of *model-based quantities and errors*. If hypotheses test main effects independently, we recommend first testing for interactions and discontinuing interpretation of main effects if significant interactions are present. To reduce erroneous conclusions and increase statistical validity, we advocate for FDR-based *multiple testing correction*. Inspection of unadjusted p-value distributions is recommended to assess model interdependence.

#### Step 5: Planning future experiments

We recommend using estimates of variance, regardless of statistical significance, to estimate sample size and power for future experiments. Within-subjects designs and exogenously defined ROIs increase signal. Rigorous QC, use of internal standards, preprocessing, filtering, and aggregation reduce noise. We recommend that all experimental procedures use *replication* and *randomization* to promote statistical validity and generalizable conclusions.

## Conclusions

This work presents a detailed accounting of each analytical step and impacts of key decisions in investigations of differential abundance, as well as an open-source workflow that can be adapted to future experiments. The workflow is specific to the goal of class comparison (i.e., detection of differentially abundant analytes). While some of the steps, such as preprocessing (Step 1) and filtering and aggregation (Step 2), are applicable to the goals of class discovery and prediction, others, such as statistical modeling and inference (Steps 3 and 4), are not.

The workflow leaves many opportunities for future research. Examples include extensions of peak picking, normalization, and feature clustering to multi-sample experiments, as well as multi-sample normalization. Future developments require the availability of high quality multi-sample datasets with known ground truth, and the development of such benchmark datasets is another important opportunity [52]. Despite the limitations, we believe that the current workflow is a step forward toward reproducible differential analysis of multi-sample MSI experiments.

## Abbreviations

ECM: extracellular matrix
MS: mass spectrometry
m/z: mass-to-charge ratio
OA: osteoarthritis
ROI: region of interest

## Acknowledgments

The authors thank Devon Kohler, Sarah Szvetecz, Anthony Wu, and Melanie Föll for technical advice and editing support. Research reported in this publication was supported by Foundation for the National Institutes of Health under award number R01AG078755. We also acknowledge the NIH for support from the following granting agencies and mechanisms: NIAT32AG000266 (to Schurman, PI:Ellerby), NIAMS R21AR084303 (MPls: Schilling, Alliston, Angel) and NIH/OD 100D030212 (PI:Angel), NIH/OD S100D038264 (PI:Schilling), and NCI R21CA240148 (PI:Angel).

## Supporting information

The following files are available free of charge.

- SupportingInformation.pdf: Includes all supporting figures, equations, and model tables (Tables S1 and S2) as well as detailed explanations and technical information.
- SupplementaryInformationTables.xlsx: Includes all non-model supporting tables (Tables S3 through S8).

