## Supplemental Information for "Statistical Principles Define an Open-Source Differential Analysis Workflow for Mass Spectrometry Imaging Experiments with Complex Designs"

#### Supporting Information

Ethan B. T. Rogers[1], Sai Srikanth Lakkimsetty[1], Kylie Ariel Bemis[1], Charles A. Schurman[2], Peggi M. Angel[3], Birgit Schilling[2], and Olga Vitek[1]

[1]Khoury College of Computer Sciences, Northeastern University, Boston MA
[2]Buck Institute for Research on Aging, Novato CA
[3]Department of Pharmacology and Immunology, Medical University of South Carolina, Charleston SC

*

### Contents

**S1 Materials and Methods** **3**

S1.1 Proposed workflow, Step 1: Data preprocessing . . . . . . . . . . . . . . . . . . . . . . . . . . 3

**Figure S1:** Simulated dataset 1: Use of multivariate segmentation for ROI segmentation decreased sensitivity of statistical tests of mean abundance between samples in Step 1 3

### S1 Materials and Methods

#### S1.1 Proposed workflow, Step 1: Data preprocessing

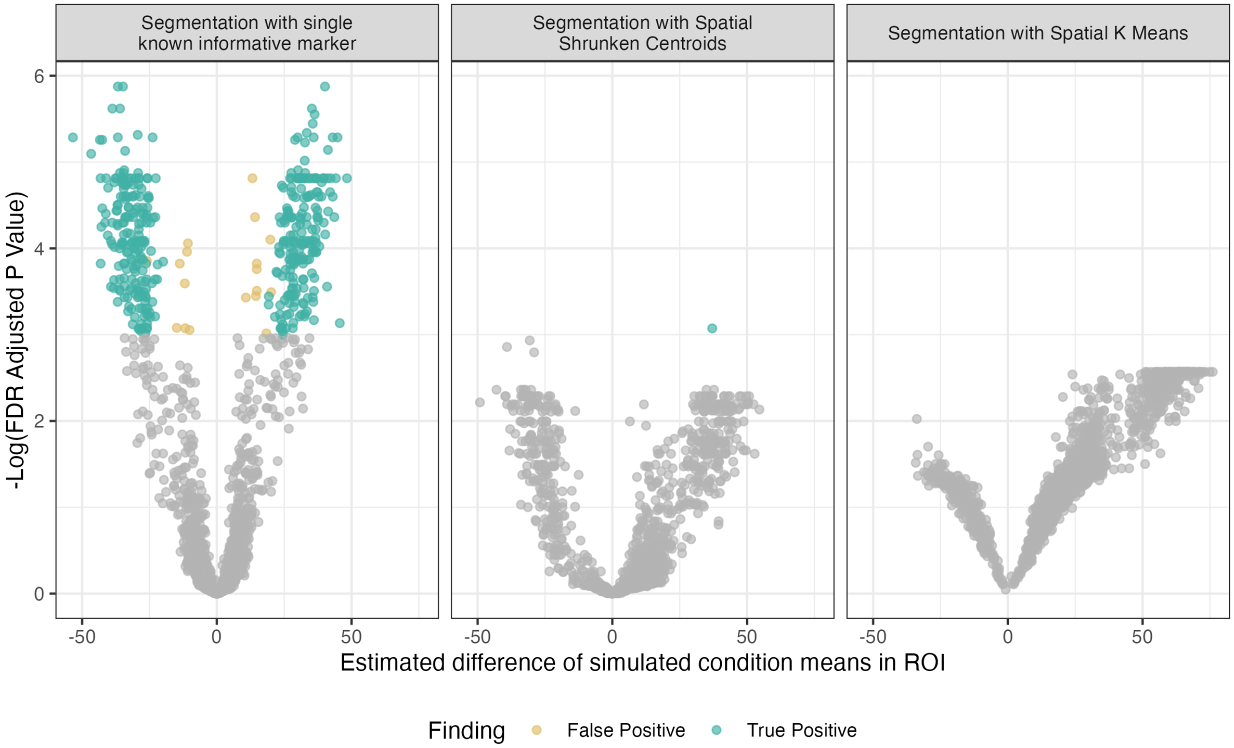

Figure S1: **Simulated dataset 1: Use of multivariate segmentation for ROI segmentation decreased sensitivity of statistical tests of mean abundance between samples in Step 1.** *Left:* Volcano plot of FDR-corrected p-values from t-tests of mean abundance between simulated conditions in the true ROI (*left*), SSC (*Middle*), and SKM (*right*). Note low sensitivity of tests performed on ROIs segmented with multivariate clustering methods.

The reduced sensitivity of statistical testing demonstrated in Figure S1 can be attributed in part to poor segmentation performance at identifying the true ROI. However, reduced sensitivity is also attributable to the confounding of information used to both segment ROIs and test for differences between them. Multivariate segmentation with SSC and SKM typically uses the entire feature set, as was the case in simulated study 1. Multivariate segmentation algorithms separate pixels into homogeneous classes based on spatial distance and shared patterns of feature intensity; as a result, corresponding classes across samples are expected to be similarly composed. This assumes that shared biology across samples produces a broadly similar set of classes, an assumption that underlies the utility of these algorithms for ROI segmentation. Under this assumption, if a feature or group of features defining a class is systematically lower in one condition, fewer of those pixels are assigned to that class, but the class mean remains largely unchanged. This selection bias attenuates detectable differences between corresponding classes across conditions, as pixels reflecting true biological variation are reassigned to other classes, absorbing what should be statistically detectable differences into small compositional shifts across segments.

```
Input: Pixel intensities X_{ijklm} from all samples, where subject i = 1...I, condition j = 1...J,
       tissue k = 1...K, pixels l = 1...n_{ijk}, and spectral features m = 1...M
Output: Clipped median-normalized pixel intensities, X_{ijklm}
// Step 1: replace zeros and clip outliers in samples
for each sample (i, j, k) do
    for each pixel l do
        Z_{ijkl} ← (X_{ijkl1}, ..., X_{ijklM});
        Z_{ijkl}[Z_{ijkl} == 0] ← NA;
        Z_{ijkl}^{0.95} ← 95th percentile of Z_{ijkl};
        Z_{ijkl}[Z_{ijkl} > Z_{ijkl}^{0.95}] ← Z_{ijkl}^{0.95};
        (X_{ijkl1}, ..., X_{ijklM}) ← Z_{ijkl};
// Step 2: Calculate scaling factor
Initialize empty 1 × IJK array SampleMedian;
for each sample (i, j, k) do
    Initialize empty 1 × n_{ijk} array PixelMedian;
    for each pixel l do
        PixelMedian_l ← median(X_{ijkl1}, ..., X_{ijklM});
    SampleMedian_{ijk} ← median(PixelMedian);
GrandMedian ← median(SampleMedian);
// Step 3: Apply scaling factor
for each sample (i, j, k) do
    for each pixel l do
        X_{ijkl}^{median} ← median(X_{ijkl1}, ..., X_{ijklM});
        for each feature m do
            X_{ijklm} ← X_{ijklm} · GrandMedian / X_{ijkl}^{median};
```

Figure S2: **OA study: Pseudocode for clipped median normalization in Step 1**

#### S1.2 Proposed workflow, Step 3: Statistical modeling

$$Y_{ijk} = \frac{1}{n_{ijk}} \sum_{l=1}^{n_{ijk}} X_{ijkl}$$

Models in Table 1 assume components ($Subject_{i(j)}$ and $\epsilon_{ijk}$) are Normally distributed and independent between measurements.

**Model 1** is a one-way ANOVA comparing conditions within a single type of tissue and equivalent to a two-sample t-test. **Model 2** is a one-way mixed-effects model comparing tissues and equivalent to a paired t test as sets of tissues belong to the same subject. Both of these models investigate the average effect of one variable within one level of the other, i.e., $Condition\ T1_j$ in Model 1 represents the systematic deviations of the measurements in condition $j$ from the overall mean of the measurements in tissue 1, and $Tissue\ C1_k$ in Model 2 represents the systematic deviations of the measurements in tissue $k$ from the overall mean of the measurements in condition 1. Therefore, these models can only investigate one hypothesis each. Model 2 reflects the fact that the measurements from a same subject are not independent, i.e. measurements from multiple tissues of a same same subject are more similar to each other than measurements from different subjects. This is reflected in Model 2 by the random term $Subject_{i(1)}$. We distinguish between-subject variance ($\sigma^2_{\text{subj}}$) and within-subject variance ($\sigma^2$) components in both Model 1 and Model 2, however only Model 2 can explicitly separate these components.

**Model 3** generalizes Models 1 and 2, using $Condition_j$ and $Tissue_k$ to represent the systematic deviations of condition $i$ and tissue $k$ from the mean of conditions and tissues respectively, and adding the interaction term $(Condition \times Tissue)_{jk}$ which reflects the fact that systematic deviations in tissues may depend on condition and vice versa. This model uses the abundances of the m/z across all the conditions and tissues to estimate the variance. This increases the precision of the estimates, as reflected in the overall degrees of freedom. Model 3 supports the same null hypotheses as the models above but can also specify more complex null hypotheses, such as Hypothesis C.

**Hypothesis A** represents the between-subjects difference of conditions in tissue 1. Only Models 1 and 3 can investigate Hypothesis A and because Model 3 uses all observations, the approximated degrees of freedom are likely to be larger than $(I-1)J$ afforded by Model 1. Higher degrees of freedom make the same test statistic less likely under the null hypothesis, increasing its significance.

**Hypothesis B** represents the within-subjects difference of tissues in condition 1 and can only be investigated by Models 2 and 3. Like for Hypothesis A, Model 3 uses all observations to estimate variances, leading to higher degrees of freedom for Hypothesis B than Model 2: $(I-1)J(K-1) > (I-1)(K-1)$. Additionally because Hypothesis A examines a within-subject effect (the difference between two tissues from the same subject, free from between-subject variance), the only source of noise to account for is residual variance. This is reflected as a smaller standard error (denominator) of Hypothesis B as compared to Hypothesis A: $\sqrt{2\frac{\hat{\sigma}^2}{I}} < \sqrt{2\frac{\hat{\sigma}^2_{subj}+\hat{\sigma}^2}{I}}$ meaning that Hypothesis B is more sensitive.

**Hypothesis C** represents the difference of differences or synergistic interaction between conditions and tissues. Only Model 3 can investigate Hypothesis C as it uses all measurements and includes an interaction term. For these reasons, we advocate for **Model 3** when modeling differential abundance in MSI experiments with complex designs, or models that use all independent variables, all measurements, and accurately partition subject and residual variance.

| Source | Type | DF | Mean squares (MS) |
|---|---|---|---|
| *Condition*, $J$ levels | Fixed | $J-1$ | $MS_{Condition} = \frac{IK \sum_{j=1}^{J} \left(\bar{Y}_{.j.} - \bar{Y}_{...}\right)^2}{J-1}$ |
| *Tissue*, $K$ levels | Fixed | $K-1$ | $MS_{Tissue} = \frac{IJ \sum_{k=1}^{K} \left(\bar{Y}_{..k} - \bar{Y}_{...}\right)^2}{K-1}$ |
| *Condition* $\times$ *Tissue*, $JK$ levels | Fixed | $(J-1)(K-1)$ | $MS_{Condition \times Tissue} = \frac{I \sum_{j=1}^{J} \sum_{k=1}^{K} \left(\bar{Y}_{.jk} - \bar{Y}_{.j.} - \bar{Y}_{..k} + \bar{Y}_{...}\right)^2}{(J-1)(K-1)}$ |
| *Subject*, $I$ levels | Random | $J(I-1)$ | $MS_{Subject} = \frac{K \sum_{j=1}^{J} \sum_{i=1}^{I} \left(\bar{Y}_{ij.} - \bar{Y}_{.j.}\right)^2}{J(I-1)}$ |
| Error | Random | $J(K-1)(I-1)$ | $MS_{Error} = \frac{\sum_{j=1}^{J} \sum_{i=1}^{I} \sum_{k=1}^{K} \left(Y_{ijk} - \bar{Y}_{i(j).} - \bar{Y}_{.jk} + \bar{Y}_{.j.}\right)^2}{J(K-1)(I-1)}$ |
| Total | | $IJK-1$ | |

Table S1: **ANOVA table of Table 1 Model 3 in Step 3, assuming balanced design with a single replicate per factor level combination.**

Table S1 details the mean squares and degrees of freedom associated with each term of Model 3 from Table 1. The mean squares characterize the variation attributable to each term in the model, on average over an infinite repetition of the experiment. Larger values indicate that more variation can be attributed to that source. Since for some models and hypotheses, specifically hypothesis A from Table1 Model 3, the degrees of freedom are not easily calculated, they are estimated using the expected mean squares. The Satterthwaite approximation is one of several methods used to calculate the degrees of freedom for estimates that contain multiple variance components, in this case subject and residual variance. For hypothesis A from Table1 Model 3 the estimate is

$$df = \frac{\left(\frac{1}{K} MS_{Subject} + (1 - \frac{1}{k}) MS_{Error}\right)^2}{\frac{\left(\frac{1}{K} MS_{Subject}\right)^2}{J(I-1)} + \frac{\left((1-\frac{1}{k}) MS_{Error}\right)^2}{J(K-1)(I-1)}}$$

The mean squares in the formula above are estimated from the data, and can be zero. For example, when subject variance is very small relative to residual variance (error), estimators may be unable to partition variances leading to an estimate of zero for $MS_{Subject}$. This may compromise the numerical stability of the Satterthwaite approximation.

| Model | Model assumptions | Null hypotheses | DF | Test Statistic |
|---|---|---|---|---|
| **Model 1:** Subset Tissue 1; One-way ANOVA comparing Conditions with pixels as replicates | $X_{ij1l} = \mu + Condition\_T1_j + \epsilon_{ij1l}$ <br> $\sum_{j=1}^{J} Condition\_T1_j = 0,$ <br> $\epsilon_{ij1l} \overset{iid^*}{\sim} \mathcal{N}(0, \psi^2_{Subj} + \psi^2)$ | **A:** Between-subject: Condition 1 vs Condition 2 in Tissue 1 <br><br> $H_0 : Condition\_T1_1 - Condition\_T1_2 = 0$ | $(IL-1)J$ | $\frac{\bar{X}_{\cdot 11\cdot} - \bar{X}_{\cdot 21\cdot}}{\sqrt{2\frac{\widehat{\psi^2_{Subj}+\psi^2}}{IL}}}$ |
| **Model 2:** Subset Condition 1; Mixed effects model comparing Tissues with pixels as replicates | $X_{i1kl} = \mu + Tissue\_C1_k + \epsilon_{i1kl}$ <br> $\sum_{k=1}^{K} Tissue\_C1_k = 0,$ <br> $\epsilon_{ijkl} \overset{iid^*}{\sim} \mathcal{N}(0, \psi^2_{Subj} + \psi^2)$ | **B:** Within-subject: Tissue 1 vs Tissue 2 in Condition 1. <br><br> $H_0 :\ Tissue\_C1_1 - Tissue\_C1_2 = 0$ | $(IL-1)K$ | $\frac{\bar{X}_{\cdot 11\cdot} - \bar{X}_{\cdot 12\cdot}}{\sqrt{2\frac{\widehat{\psi^2_{Subj}+\psi^2}}{IL}}}$ |
| **Model 3**: Mixed effects model for all Conditions and Tissues with pixels as replicates | $X_{ijkl} = \mu + Subject_{i(j)} + Condition_j +$ $Tissue_k + (Condition \times Tissue)_{jk} + \epsilon_{ijk}$ <br> $\sum_{j=1}^{J} Condition_j = 0, \sum_{k=1}^{K} Tissue_k = 0,$ <br> $\sum_{j=1}^{J} (Condition \times Tissue)_{jk} = 0,$ <br> $\sum_{k=1}^{K} (Condition \times Tissue)_{jk} = 0,$ <br> $Subject_{i(j)} \overset{iid}{\sim} \mathcal{N}(0, \psi^2_{subj}),\quad \epsilon_{ijk} \overset{iid^*}{\sim} \mathcal{N}(0, \psi^2)$ | **A:** Between-subject: Condition 1 vs Condition 2 in Tissue 1 <br><br> $H_0 :\ Condition_1 + (Condition \times Tissue)_{11} - [Condition_2 + (Condition \times Tissue)_{21}] = 0$ | Satterthwaite approximation, see below | $\frac{\bar{X}_{\cdot 11\cdot} - \bar{X}_{\cdot 21\cdot}}{\sqrt{2\frac{L\widehat{\psi^2_{Subj}+\psi^2}}{IL}}}$ |
| | | **B:** Within-subject: Tissue 1 vs Tissue 2 in Condition 1. <br><br> $H_0 :\ Tissue_1 + (Condition \times Tissue)_{11} - [Tissue_2 + (Condition \times Tissue)_{12}] = 0$ | $(I-1)J(K-1) + IJK(L-1)$ | $\frac{\bar{X}_{\cdot 11\cdot} - \bar{X}_{\cdot 12\cdot}}{\sqrt{2\frac{\hat{\psi}^2}{IL}}}$ |
| | | **C:** Difference of differences: Differences between Conditions 1 and 2 in Tissue 1 to that of Tissue 2. <br><br> $H_0 :\ [(Condition \times Tissue)_{11} - (Condition \times Tissue)_{12}] - [(Condition \times Tissue)_{21} - (Condition \times Tissue)_{22}] = 0$ | $(I-1)J(K-1) + IJK(L-1)$ | $\frac{\bar{X}_{\cdot 11\cdot} - \bar{X}_{\cdot 12\cdot} - \bar{X}_{\cdot 21\cdot} - \bar{X}_{\cdot 22\cdot}}{\sqrt{4\frac{\hat{\psi}^2}{IL}}}$ |

Table S2: **Statistical models in Step 3 using pixels as replicates for experiments Figure 11 with balanced designs and no missing values.** The models assume balanced designs and no missing values. $X_{ijkl}$ denotes the intensity of pixel $l = 1, \ldots, L$, of the feature in the ROI of a sample of subject $i = 1, \ldots, I$, condition $j = 1, \ldots, J$, and tissue $k = 1, \ldots, K$. Model terms with zero-sum constraints are fixed effects, and are the parameters of interest. Model terms that follow a Normal distribution are random effects. $\bar{X}$ is the intensity averaged over the index indicated with a dot. $\hat{\psi}$ is the data-derived estimate of variation, artificially decreased by neighboring pixel autocorrelation. *Violation of assumption: not independent.

Table S2 details the same models presented in Table 1 but specifying pixel intensity, $X_{ijkl}$, as their response variable and using pixels as replicates. Using pixels as replicates adds thousands of "replicates", reflected by the inclusion of $l$ in the model specifications and $L$, the total number of pixels, in the degrees of freedom and test statistics. However these pixels are not independent replicates, as individual pixel intensity is spatially correlated with neighboring pixels. Adding pixels increases the degrees of freedom for every comparison. Additionally, standard errors (denominator of the test statistic, e.g., $\sqrt{2\frac{\hat{\psi}^2}{IL}}$ for hypothesis B from Model 3 in Table S2 ) are systematically lower as variance ($\hat{\psi}^2$) is spread over subjects *and* pixels ($IL$). Pixel autocorrelation further reduces the estimates of variances. We differentiate these systematically low estimates of variation using $\psi^2$ and $\psi^2_{Subj}$ instead of $\sigma^2$ and $\sigma^2_{Subj}$. Additionally, unlike Table 1 Model 2 specifying mean pixel intensity as the response variable, Table S2 Model 2 specifying pixels as replicates is not a paired design, as tissues from the same subject are not the same size and do not have paired pixels.

For hypothesis A from Table S2 Model 3, degrees of freedom are estimated using the Satterthwaite approximation.

$$\frac{\left(\frac{1}{K}MS_{Subject} + (1-\frac{1}{k})MS_{Error}\right)^2}{\frac{\left(\frac{1}{K}MS_{Subject}\right)^2}{J(I-1)} + \frac{\left((1-\frac{1}{k})MS_{Error}\right)^2}{J(K-1)(I-1)+IJK(L-1)}}$$

Using pixels as replicates inflates the degrees of freedom for each mean square component, reducing the size of the denominator and increasing the approximation. A larger Satterthwaite approximation of degrees of freedom contributes to the increased, though erroneous, sensitivity of Model 3 using pixels as replicates.

#### S1.3 Proposed workflow, Step 4: Statistical inference

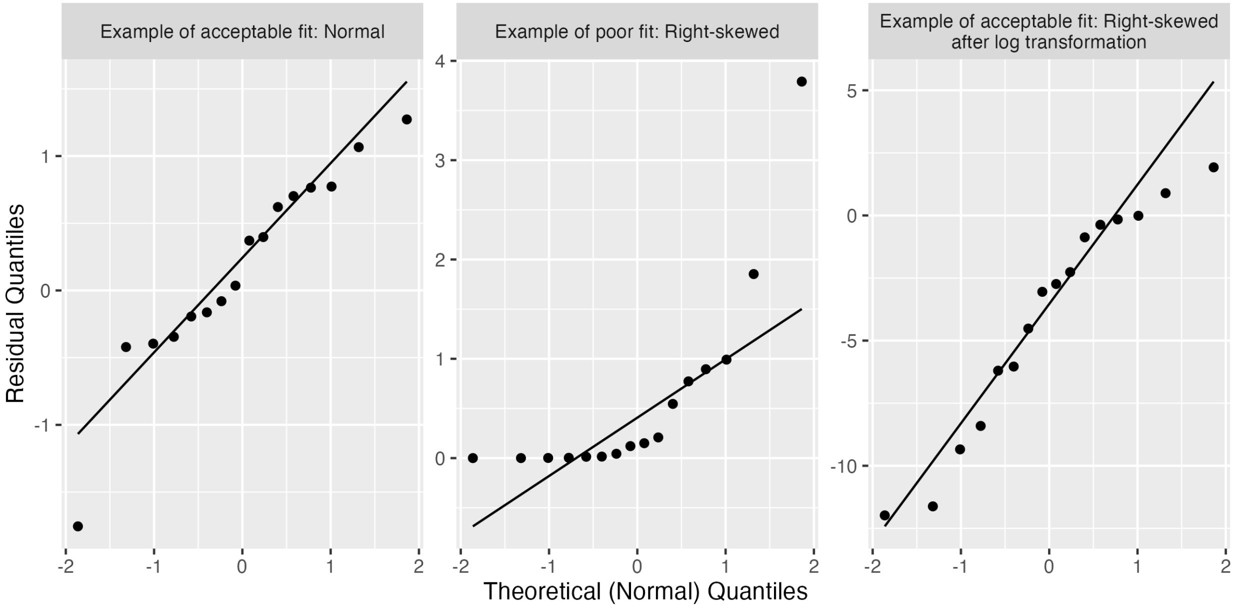

Figure S3: **Examples of residual quantile-quantile plots demonstrating fit quality.**

Non-linear quantile-quantile plots indicate deviance from the assumption of normality. Figure S3 offers three examples of different fits. Normally distributed residuals should generally follow the reference line (Figure S3 *left*). Some deviation is expected but serious deviations, such as for right skewed residuals (Figure S3 *middle*), indicate poor fit due to the model's assumption of normally distributed residuals not being fulfilled. Some types of deviations, such as right skew, can be corrected with transformations (Figure S3 *right*). The informative value of diagnostic plots diminishes with smaller sample sizes, as patters may be harder to spot.

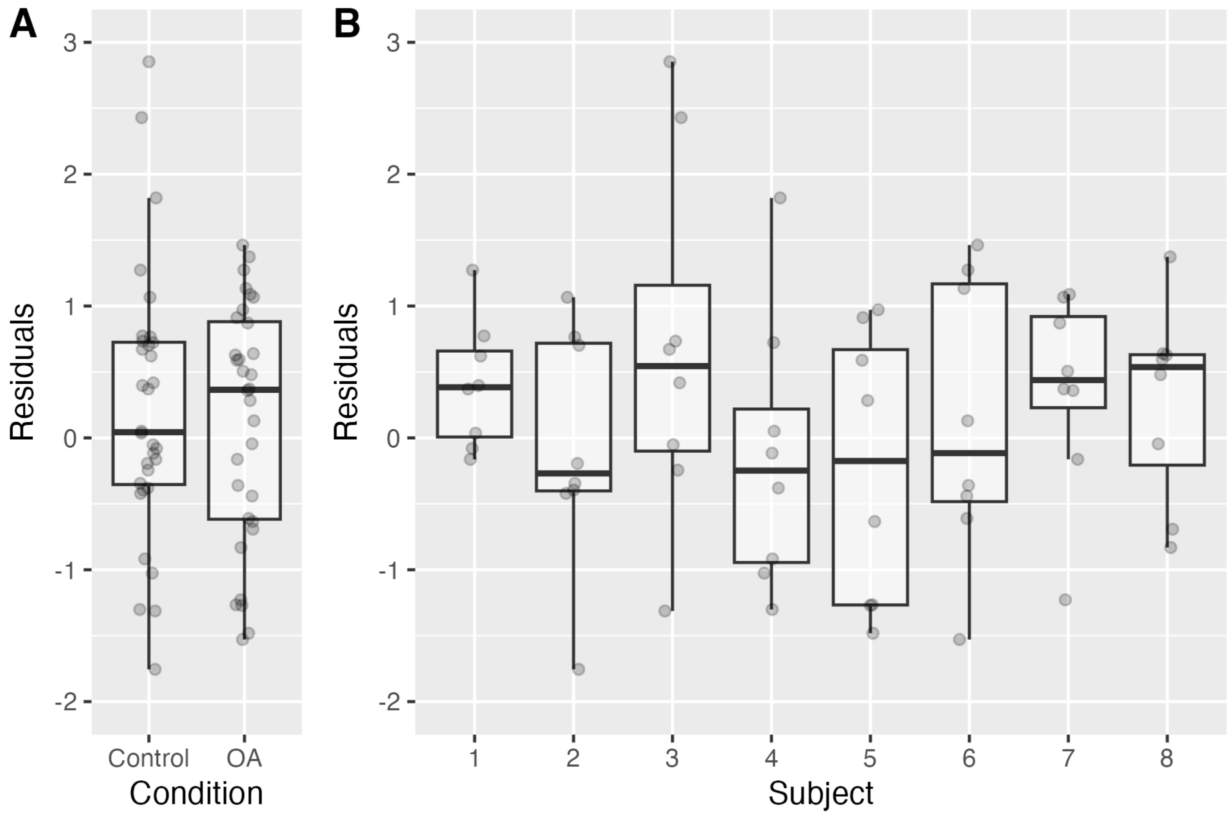

Figure S4: **Examples of residual box plots demonstrating acceptable equality of variance across conditions and tissues. A:** Simulated Normal residuals by condition. **B:** Simulated Normal residuals by subject with 8 measurements per subject.

Linear models also assume equal variation of residuals within each group, specifically between levels of independent variables (e.g., between conditions in Figure S4A) and random factors (between subjects in Figure S4B). Box plots can be used to determine if the assumption of constant variance is met with categorical independent variables. As with quantile-quantile plots, assessments of constant variance are less meaningful at smaller sample sizes. For this reason, we did not assess this assumption in the OA dataset.
